# Evolution of aberrant brain-wide spatiotemporal dynamics of resting-state networks in a Huntington’s disease mouse model

**DOI:** 10.1101/2023.11.20.567961

**Authors:** Tamara Vasilkovska, Marlies Verschuuren, Dorian Pustina, Monica van den Berg, Johan van Audekerke, Isabel Pintelon, Roger Cachope, Winnok H. De Vos, Annemie Van der Linden, Mohit H. Adhikari, Marleen Verhoye

## Abstract

Distinct resting-state networks (RSNs) are differentially altered in the course of Huntington’s disease (HD). However, these RSN changes are depicted using traditional functional connectivity analyses which ignore the dynamic brain states that constitute these RSNs and their time-dependent relationship. Dynamic states are represented by recurring spatiotemporal patterns of propagating cortical and subcortical brain activity observed in low-frequency BOLD fluctuations, called quasi-periodic patterns (QPPs). In this study, we used resting-state fMRI to investigate QPPs in the zQ175DN mouse model of HD at 3, 6 and 10 months of age. We identified age- and genotype-specific short (3s) QPPs, representative of the lateral cortical network (LCN) and the default mode-like network (DMLN), and a long (10s) QPP, the homolog of the human primary QPP, exhibiting the propagation of activity between the LCN and the DMLN. Hyperactivity was present in the caudate putamen and the somatosensory cortex in zQ175DN mice at 3 and 6 months of age. Moreover, DMLN-wide reduction in activation was observed at all ages, where at 6 and 10 months of age the reduced activity gradually advanced into a breakdown of the LCN-to-DMLN propagation. We then investigated the relationship in the timing of peak activity of six brain regions involved in the long QPPs and found that the retrosplenial cortex, a transmodal region which orchestrates multisensory integration, has a premature peak of BOLD activation in zQ175DN mice at 6 months of age, as compared to age-matched controls. Irrespective of either LCN or DMLN activation, this resulted in an asynchrony of the retrosplenial cortex in the peak timing relationships relative to other regions during the long (10s) QPPs. Finally, the normative, age-dependent, wild-type QPPs were significantly decreased in occurrence in the zQ175DN group at each age, indicating the presence of phenotypically-driven LCN and DMLN states as captured with QPPs. As BOLD-dependent variations result from neurovascular coupling, we assessed mutant huntingtin (mHTT) deposition in astrocytes and pericytes, known components of the neurogliovascular unit. These analyses showed increased cell-type dependent deposition starting at 6 months in the caudate putamen, somatosensory and motor cortex, regions that are prominently involved in HD pathology as seen in humans. Our findings provide meaningful insights into the development and progression of altered functional brain dynamics in this HD model, opening potential new avenues for its application in clinical HD research.

**SUMMARY:** Huntington’s disease (HD) is marked by irreversible loss of neuronal function for which currently no availability for disease-modifying treatment exists. Advances in the understanding of disease progression can aid biomarker development, which in turn can accelerate therapeutic discovery. We characterized the progression of altered dynamics of whole-brain network states in the zQ175DN mouse model of HD using a dynamic functional connectivity (FC) approach to resting-state fMRI and identified quasi-periodic patterns (QPPs) of brain activity constituting the most prominent resting-state networks. The occurrence of the normative QPPs, as observed in healthy controls, was reduced in the HD model as the phenotype progressed. This uncovered progressive cessation of synchronous brain activity with phenotypic progression, which is not observed with the conventional static FC approaches. This work opens new avenues in assessing the dynamics of whole brain states, through QPPs, in clinical HD research.

## INTRODUCTION

Huntington’s disease (HD) is the most prevalent monogenic neurodegenerative disorder and genetic dementia, characterized by the progressive loss of cognitive and motor function^1^. Despite the known genetic background^2,3^, there is insufficient understanding of the progression of the functional phenotype of HD. MRI neuroimaging has been the cornerstone in unraveling the brain’s structural and functional alterations across HD progression due to its noninvasive nature, thus enabling the discovery of novel biomarkers. Currently, detection of striatal brain loss using volumetric MRI is by far the most sensitive and earliest biomarker, manifesting even 20 years prior to HD clinical motor diagnosis^2,4–6^. However, even with this method, brain loss has already occurred, which is both irreversible and can progressively worsen over time. There is a critical need for novel early detection methods prior to irreversible brain atrophy.

A promising but underexplored MRI technique in HD is resting-state functional MRI (rsfMRI)^7^. rsfMRI is used to investigate changes in large-scale network connectivity, revealing the brain’s functional architecture, which is organized in resting-state networks (RSNs)^8–10^. Mapping functional connectivity (FC) in both the healthy brain and in neurodegenerative diseases (ND)^11,12^, including HD, has shown dysfunction in several RSNs^7^ before and after clinical motor diagnosis in people with HD (PwHD)^13–15^. However, these findings represent static FC, which assumes a constant correlation between a pair of brain regions and flattens the changes in time. This provides only snapshots of the brain, while the activity states of the brain are highly dynamic. Neglecting the dynamics of brain activity leads to an incomplete picture of the disease’s pathological effect on the brain, which significantly limits fundamental understanding, identification of biomarkers, and the development of novel therapies.

To address this issue, novel methods are utilized to account for time-varying properties and identify dynamic brain states that underlie the RSNs^16,17^. These methods elucidate time-evolving activity across whole brain spatial patterns, which have led to the discovery of quasi-periodic patterns (QPPs) that encompass cyclic brain activity^18^. The most prominent human QPP, the primary QPP, represents the anti-correlated activity of the two most dominant RSNs, the default-mode network (DMN) and the task-positive network (TPN)^19^. Alterations in these dynamic brain states, as revealed by QPPs, have also been shown in several neuropsychiatric disorders^20,21^. While the presence of these dynamic brain patterns has been detected in the healthy human brain, they have not been evaluated in any humans presenting with ND. Recently, our lab has demonstrated the potential of QPPs to assess disease-driven alterations in rodent models of ND, specifically Alzheimer’s’ disease^22,23^.

Here, we assessed the evolution of dynamic brain-wide network changes across disease-like progression in the zQ175DN knock-in (KI) heterozygous (HET) mouse model of HD. We hypothesized that QPPs can be used to detect functional alterations in whole brain activity in this HD mouse model that will give novel insights into the disease-like phenotype and uncover changes at ages before brain atrophy is observed.

We found a progressive cessation of synchronous network activity in the mouse default mode-like network (DMLN), whose gradual reduction in activation led to the absence of the primary QPP rodent homolog. These results were further validated in histological staining for cells of the neurovascular unit within the brain. Moreover, within the QPP the retrosplenial cortex, a central hub of the DMLN, showed asynchrony in the peak timing relationship with other regions irrespective of the network state. This indicates a potential cause of the breakdown of DMLN activity, concurrent with the putative onset of motor deficits in this mouse model. Finally, using QPP projection, we found that the normative QPPs that occur in healthy controls were consistently and progressively reduced along phenotypic progression in this HD model. The initial reduced states were present even before brain volume loss, suggesting that this method holds great potential for its application in clinical HD research. Altogether, our results provide a significant understanding of the progression of brain circuitry alterations in this HD mouse model, which encourages exploring if similar differences can be detected in PwHD, even before brain atrophy and clinical motor diagnosis occur.

## MATERIALS & METHODS

### In-vivo resting-state fMRI study

#### Animals

A total of 40 mice, male, age-matched, HET (*n* = 20) zQ175DN KI and WT (*n* = 20) littermates (C57BL/6J background, CHDI-81003019, JAX stock #029928) were obtained from the Jackson Laboratory (Bar Harbor, ME, USA). Animals were single-housed in individually ventilated cages with food and water *ad libitum* and continuous monitoring for temperature and humidity under a 12h light/dark cycle. The animals were kept in the animal facility for at least a week to acclimatize to the current conditions before the experimental procedures. All experiments and handling were done in accordance with the EU legislation regulations (EU directive 2010/63/EU) and were approved by the Ethical Committee for Animal Testing UAntwerp (ECD # 2017-09).

The zQ175DN KI (DN: delta neo - without a floxed neomycin cassette) mouse model has the human HTT exon 1 substitute for the mouse *Htt* exon 1 including a ∼180-220 CAG repeats long tract. This model is a modified version of the zQ175 KI^24^ where the congenic C57BL6J is used as the strain background^25^. In this animal model, the initial motor deficits are detected by 6 months of age, marked as an onset of phenoconversion^26^. The HET form of zQ175DN has a slow progression in terms of neuropathology, reflected in the increase of mutant huntingtin (mHTT) protein aggregation from 3 until 12 months, initially appearing in the striatum at 3 and later in the cortex at 8 months of age^27^. A longitudinal rsfMRI study was performed in both the zQ175DN HET and WTs^28^ at ages of 3, 6 and 10 months, following phenotypic progression.

#### Image acquisition

MRI scans were acquired on a 9.4 T Biospec system with a four-element receive-only mouse head cryoprobe coil (Bruker Biospin MRI, Ettlingen, Germany) and a volume resonator for transmission. Prior to the whole brain rsfMRI scan, a B_0_ field map was acquired to measure the magnetic field inhomogeneities after which local shimming was performed within an ellipsoid volume, covering the middle portion of the brain. The dynamic blood oxygen level-dependent (BOLD) resting-state signals were measured with a T2*-weighted single-shot Echo-Planar Imaging (EPI) sequence (field of view (FOV) (27×21) mm^2^, matrix dimensions (MD) [90×70], 12 horizontal slices of 0.4mm, voxel dimensions (300×300×400) µm^3^, flip angle 60°, TR 500ms, TE 15ms, 1200 repetitions). After the rsfMRI scan was acquired, a 3D anatomical scan was obtained using the 3D Rapid Acquisition with Refocused Echoes (RARE) sequence with FOV (20×20×10) mm^3^, MD [256×256×128], voxel dimensions (78×78×78) µm^3^, TR 1800ms, TE 42ms.

During rsfMRI scans, mice were initially anaesthetized with 2% isoflurane (Isoflo®, Abbot Laboratories Ltd., USA) in a mixture of 200ml/min O_2_ and 400ml/min N_2_. After the animal was positioned in the scanner, a subcutaneous bolus injection of medetomidine (0.075 mg/kg; Domitor, Pfizer, Karlsruhe, Germany) was applied followed by a gradual decrease of isoflurane to 0.5% over the course of 30 minutes which was kept at this level throughout the whole experiment. Meanwhile, a continuous s.c. infusion of medetomidine (0.15 mg/kg/h), starting 10 min post bolus medetomidine injection, was applied in combination with the isoflurane, an established protocol for rsfMRI in rodents. Throughout the duration of the experiment, all physiological parameters (breathing rate, heart rate, O_2_ saturation, and body temperature) were kept under normal conditions.

The experiments were continuously alternated between WT and zQ175DN HET for all ages. Mice were randomized for each acquisition at each age. All data were collected by using animal ID and blinding was only applied during the allocation of animals. Prior to the experiment, weight measurements obstructed potential blinding due to genotype-driven weight differences. One animal from each group was mis-genotyped and one WT deceased. Finally, a total of 37 animals (WT (*n* = 18) and zQ175DN HET (*n* = 19)) were used for further analysis.

#### Image preprocessing

rsfMRI repetitions for each session were realigned to the first image with a 6-parameter rigid-body spatial transformation estimated with the least-squares approach (limited head movement with displacement <0.4mm in either direction). Based on the obtained realignment parameters, motion vector regression was applied. Next, a study-based template was generated. To create an unbiased template, we used the individual 3D RARE images from 1/2 of each group from each time point. The rsfMRI data were spatially normalized to their respective subject 3D RARE image. The 3D RARE images were normalized to the common study template. Spatial transformation parameters were also estimated between the study-based template and an in-house C57BL/6 mouse brain atlas. All the rsfMRI data were spatially normalized to the in-house C57BL/6 atlas, combining all estimated (rigid, affine, and non-linear) transformation parameters: (1) rsfMRI to 3D RARE, (2) 3D RARE to common study template, and (3) common study template to in-house C57BL/6 atlas. In-plane smoothing was applied using a Gaussian kernel with full width at half maximum of twice the voxel size followed by motion vector regression based on the parameters generated during realignment. These preprocessing steps were performed using Statistical Parametric Mapping (SPM) using SPM12 (Wellcome Centre for Human Neuroimaging, London, UK). Using a whole-brain mask, images were further filtered (0.01-0.2Hz) with a Butterworth band-pass filter where five repetitions from both the beginning and the end of the image series were removed before and after filtering to eliminate transient effects. Finally, quadratic detrending (QDT), voxel-wise global signal regression (GSR), and normalization to unit variance were applied. Template creation was done using Advanced Normalization Tools (ANTs).

#### QPP analysis

QPP extraction was done at the group level for each age separately, since previously, we have shown an age-dependent FC change in this HD model^29^. First, the rsfMRI scans from all subjects were concatenated in one image series for WT and zQ175DN HET. A spatiotemporal pattern finding algorithm^30^ was then applied, which extracts whole-brain recurring patterns of BOLD activity in a user-defined temporal window. Next, the optimal time window of the sliding template was estimated^31^, which resulted in two window sizes of 6TR (3s) and 20TR (10s). Consecutive BOLD image frames with the predefined window length from a random starting frame in the image series are taken as an initial template. Its similarity with the entire image series is then stepwise calculated by estimating its Pearson correlation with image-series segments of the same window size and progressively shifting it by 1 TR. Image frames with local maxima of the resulting sliding template correlation (STC) that surpass a correlation threshold of 0.2 were then voxel-wise averaged to form an updated template. The process was repeated using the updated template until the correlation coefficient between templates from consecutive iterations exceeded 0.99 at which point the template was referred to as a QPP. The entire procedure was reiterated for 200 random starting frames from the image series resulting in 200 short (6 TR) and 200 long (20 TR) QPPs.

For both short and long QPPs, in order to identify (dis)similar QPPs within the 200 QPPs, hierarchical clustering was performed on the Euclidean norm of both spatial and temporal domain distances. Robust clusters were determined based on several criteria: a minimum size of 5% of the total number of QPPs, a minimum of 5% occurrence from total occurrences, and the majority of QPPs within each cluster must occur at least in 80% and 50% of subjects for short and long QPPs, respectively. The representative QPP (rQPP) from each cluster was obtained as the QPP with the highest total correlation (summed across occurrences) with the image series.

To assess the recurring property of a QPP, occurrences of each rQPP for each subject were extracted based on the rQPP’s STCs. The occurrence rate of a QPP is defined as the number of occurrences per minute (Occurrences/min).

At each age, short and long rQPPs from both WT and zQ175DN HET groups were projected onto the image series of the other group to identify their representative spatial pattern and occurrences in the other group. Projection involved the calculation of the STC of the rQPP from one group with the image series of the other group and finding its local maxima exceeding the threshold of 0.2 that yielded the projected occurrence rates. Voxel-wise averaging across projected occurrences resulted in the projected pattern in the other group.

#### QPP regression

To understand the contribution of QPPs to FC, linear regression of the rQPPs was performed within each group’s image series^22^. The rQPP was convolved with its own STCs to create that QPP’s image series. Further, this QPP image series was voxel-wise regressed from the original image series, thus removing the QPP’s influence on the BOLD signal. The residual scans were used to further assess the FC without the contribution of that QPP. The QPP regression was performed for short rQPPs in both WT and zQ175DN HET groups.

#### FC analysis

Static FC between selected regions of interest (ROIs) that pertain to four different networks: the DMLN, associative cortical network (ACN), lateral cortical network (LCN) and the Subcortical Network (SuCN) was calculated. We selected 26 unilateral ROIs (both left (L) and right (R) hemispheres for each region) from an in-house C57BL6 mouse atlas^32^ that represent the main hubs of several relevant large-scale networks^33,34^. The abbreviations for each region are presented in Table 1. For each subject and each region, we extracted the time series of the region-averaged BOLD signal. Pearson correlation coefficients were calculated between the BOLD signal time series of each pair of regions. These correlations were Fisher z-transformed, thus obtaining subject-wise FC matrices.

**Table 1.**
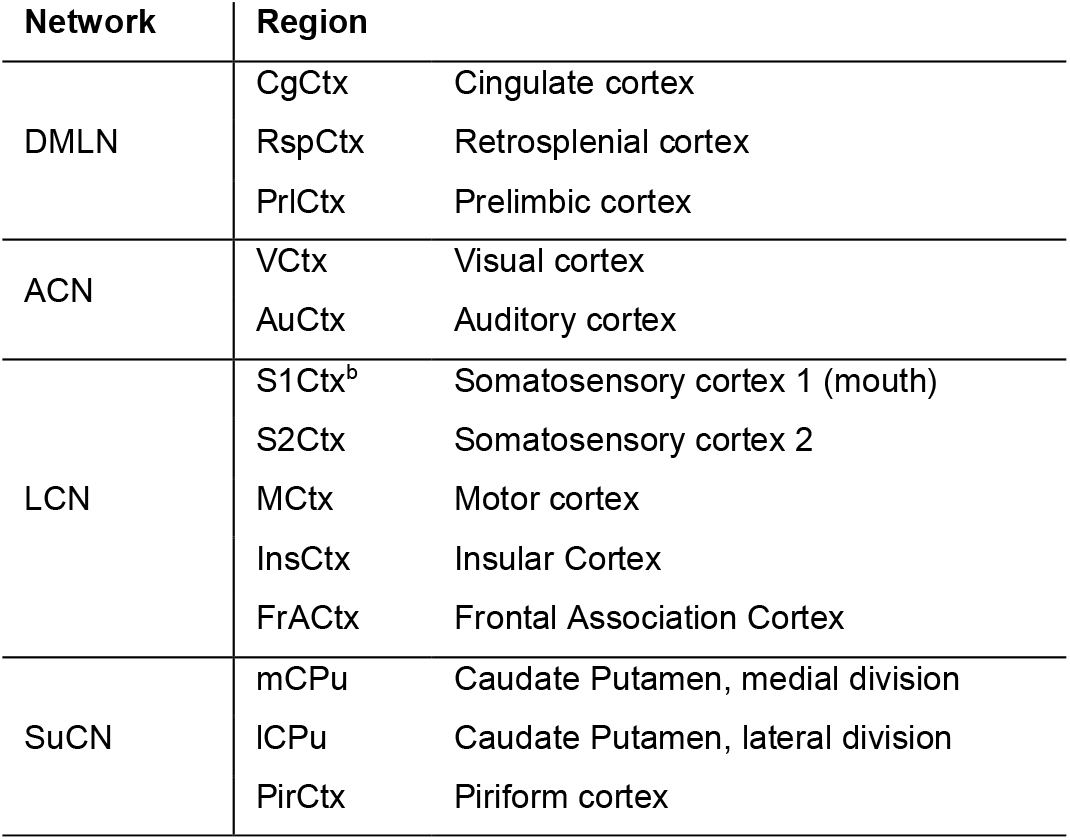
Brain regions and Network abbreviations.

#### Regional peak timing relationship within a long rQPP

To understand the propagation of coordinated activity in key regions involved in the long QPPs and to unravel potential driving mechanisms, an adapted method for extraction of peak timing of regional (de)activation was applied^19^. The first step was the identification of significantly (de)activated voxels within the whole duration of 10s of the long rQPP. To obtain these, we first constructed a null spatiotemporal pattern by voxel-wise averaging across n, randomly selected, non-overlapping segments that do not overlap with segments corresponding to occurrences of a QPP of interest, where n is the number of occurrences of that QPP within the image series. The number of random segments from a single subject corresponds to the number of times the QPP occurs in that subject. This process is repeated 50 times resulting in 50 null QPPs.

Next, the root sum square (RSS) of the voxel-wise BOLD signal was calculated for each QPP of interest and the corresponding null QPPs. Thus, for each voxel, its BOLD signal for each of the 20-time frames of the QPP was squared, summed across all frames and then a square root of this sum was taken. This resulted, for the 50 null QPPs, in 50 x 7688 RSS values, where N is the number of voxels in the brain mask we considered. Voxels whose RSS values in each long rQPP exceeded the 95 percentile of the null QPP RSS values were identified as significant and used for further analysis. These voxels were assigned to six regions: Caudate putamen (CPu), medial prefrontal cortex (mPFC), motor cortex (MCtx), Somatosensory cortex 1 (S1Ctx), retrosplenial cortex (RspCtx) and visual cortex (VCtx). Each voxel shows two peaks of absolute values of its BOLD signal during every occurrence segment of the QPP (e.g., in the long (10s) rQPP, for most voxels in the S1Ctx, there is an activation (>0) peak during the first 10 frames of the QPP segment (LCN) and a second peak of deactivation (< 0) during the last 10 frames of the QPP segment (DMLN)). For each voxel within the QPP, timings for both the first and second peaks are extracted for all occurrences of the long QPP. All timings for each active voxel were averaged across the occurrences within each group.

#### Statistical analysis

To visually inspect the significant voxel-wise (de)activation that constitutes an rQPP, one-sample T-statistic maps were calculated based on the BOLD signal across all occurrences of that QPP in the image series (One-sample T-test, False Discovery Rate (FDR) corrected, p < 0.05, cluster size (k) ≥ 10). Between-group differences in spatial voxel activations, occurrence rates and FC were cross-sectionally compared for each age (Two-sample T-test, FDR corrected, p < 0.05, k ≥ 10). For a visual representation of the anatomy of prominent QPPs, the T-statistics were transformed and upsampled to the high-resolution Australian mouse brain MRI anatomical atlas^35^ using transformation parameters calculated between the C57BL6 mouse atlas and the Australian mouse brain atlas. These visualizations were produced with MRIcroGL (McCausland Center for Brain Imaging, University of South Carolina, USA).

For the regional peak timings in the long rQPP, the mean timing for each peak across all activated voxels was first compared between genotypes (Two-sample T-test, FDR corrected, p < 0.05). Separately for the first and second peak, the mean peak timing difference between every pair of regions, within a genotype (Two-sample T-test, FDR, p < 0.05) was assessed. The final comparison identified region-pairs whose peak timings relationship was significantly altered between genotypic groups (Two-sample T-test, FDR corrected, p < 0.05). To compensate for the difference in the number of significantly activated voxels of different regions, we calculated the difference between the peak timing of each voxel of the smaller region with the mean (across voxels) peak timing of the bigger region for the last comparison. Multiple comparison corrections were applied for both metrics (timings of two peaks) and over all regions or region-pairs (FDR, p<0.05). All analysis steps and statistics were performed with MATLAB R2021a (Mathworks, Natick, MA).

### Ex-vivo immunofluorescence (IF) study

#### Brain preparation

To assess changes in astrocytes and pericytes in order to understand impairments in neurovascular coupling, an IF study was performed on 12 age-matched, male WT (n = 6) and zQ175DN HET (n = 6) mice at each of the 3, 6, 8 and 12 months of age (48 animals in total). Cardiac perfusion was performed with PBS solution and 4% paraformaldehyde solution (PFA). Extracted brains were post-fixed in 4% PFA at 4°C for 24h, followed by 30% sucrose at 4°C for 72h. Brains were cryo-embedded in Tissue-Tek® O.C.T. Compound (Sakura, Cat. # 4583) with a precision cryo-embedding system and stored at -80°C. For sectioning, brains were embedded in an O.C.T.-embedding medium. At Bregma lateral 2.52, 0.96 and 0.12mm, sagittal sections of 30 µm were obtained using a Leica CM1950 cryomicrotome (Leica BioSystems, Belgium), thaw mounted on VWR Superfrost Plus micro slides (VWR, Leuven, Belgium) and dried for 2 h at 37 °C.

#### IF staining

Sections were first acclimatized at room temperature (RT) and further rehydrated in PBS (Life Technologies, 7001-36) for 5 min. Next, antigen retrieval was done by incubation in citrate buffer pH 6.0 for 30 min at 80°C. After washing three times with PBS, permeabilization and blocking of endogenous proteins were done with 0.5% Triton X-100 and 10% normal donkey serum (NDS, Jackson Immunoresearch, 017-000-121) in PBS for 30 min, followed by donkey anti-mouse IgG Fab fragment (1:40 in PBS, Jackson Immunoresearch, 715-006-151), blocking for 1h at RT. Two stainings were performed where the following primary antibodies were applied in the first staining (1) 2B4 (1:200, Sigma Eldrich, Cat. # MAB5492) and CD13 (1:100, R&D Systems, Cat. # AF2335) diluted in PBS + 5% NDS and in the second staining (2) 2B4 (1:200, Sigma Eldrich, Cat. # MAB5492), GFAP (1:1000, Abcam, Cat. # ab4674) and VEGF (1:100, Thermo Fisher, Cat. # MA5-32038) were diluted in 0.1% Triton X-100 and 1% NDS in PBS and incubated overnight at RT. After washing with PBS, the fluorescent-conjugated secondary antibodies were applied: (1) donkey anti-mouse Cy5 (1:50, Jackson Immunoresearch, Cat. # 715-175-151) for 2B4 and donkey anti-goat AF555 (1:200, Thermo Fisher, Cat. # A21432) for CD13 and (2) donkey anti-chicken Cy3 (1:500, Jackson Immunoresearch, Cat. # 703-166-155), donkey anti-mouse Cy5 (1:50, Jackson Immunoresearch, Cat. # 715-175-151) and donkey anti-rabbit AF488 (1:200, Jackson Immunoresearch, 711-545-152) for VEGF with PBS for 1h on RT. In the first staining, an additional subsequent 2h incubation with Lectin-Dylight 488 (1:25, Vector Laboratories, Cat. # DL-1174). After washing, in both stainings, sections were covered with a coverslip with antifade mounting medium containing DAPI (VECTASHIELD® Antifade Mounting Medium with DAPI Cat. # H-1500) and stored at 4°C.

#### IF image acquisition

Confocal images of the stained sections were acquired for 8 brain ROIs: medial (m) CPu (lateral 0.96mm), lateral (l) CPu (lateral 2.52mm), S1Ctx, MCtx 1, MCtx 2, Cingulate Cortex (CgCtx), Piriform Cortex (PiriCtx) and Insular Cortex (InsCtx) with a Perkin Elmer Ultraview Vox dual spinning disk confocal microscope, mounted on a Nikon Ti body using a 40x dry objective (numerical aperture 0.95). Lasers with wavelengths 405 nm, 488 nm, 561nm and 640nm were used in combination with a quadruple dichroic and 445/60 - 525/50 - 615/70 - 705/90 emission filters. Detection was done on a Hamamatsu C9100-50 camera. Image acquisition was done using the Volocity software. Per animal and brain region, 5 images were acquired in 30 axial positions separated by a 1 μm spacing and a pixel resolution of 0.1786 µm.

#### IF image processing and analysis

A macro script was written for FIJI image analysis^36^ and is available on github (https://github.com/DeVosLab/Huntington). This script is built on modules that were already integrated with similar image analysis pipelines previously developed^37–39^. In brief, 30µm z-stacks were divided into maximum projections of 5µm substacks, with 2µm in between each substack. In each projection, the DAPI channel was used to detect nuclei employing the Stardist-trained convolutional network^40^ after background correction and smoothing (gaussian filter with a radius of 8 pixels) to suppress contrast variations. Identified nuclei were then filtered based on user-defined area and circularity limits. To define a cell region, the nuclear region was concentrically dilated by 3.2µm. Due to region-dependent differences in signal distribution (e.g. 2B4 signal, Fig. S10), we opted for region specific analysis settings that were applied for all animals and time points within the specified region. 2B4-positive huntingtin spots were segmented based on a user-defined threshold after background subtraction and multi-scale Laplacian spot enhancement. Spots were further filtered based on user-defined area and circularity filters. Non-specific 2B4 staining in blood vessels was excluded from this analysis. Cells that contained at least one spot were identified as 2B4+ cells. 2B4 spot coverage was calculated as the sum of the area of all spots in a cell divided by the projected cell area. To identify blood vessels and CD13+ vascular regions in the maximum projections, a multi-scale tubeness filter^41,42^ and a user-defined threshold were used on both the lectin and CD13 channels, respectively. CD13 coverage on blood vessels was defined as the area of the overlapping CD13+ regions and lectin+ regions divided by the total blood vessel area (lectin+) in a FOV. To quantify the area of GFAP+ regions, a Gaussian filter (7-pixel kernel radius) was applied to the maximum projections after which a low user-defined threshold was applied to detect all GFAP+ regions in the image.

After processing, data analysis was performed in R^43^. Cells that had a nuclear overlap with CD13+/GFAP+ pixels greater than 50% were assigned as CD13+ or GFAP+ cells respectively. All single-cell data from substack projections were pooled for every image to obtain the final cell counts. Other variables (*e.g.,* spot coverage) were averaged over all substack projections for each image.

#### IF statistical analysis

Image-based outlier detection was applied using the median ± 3 x median absolute deviation as outer limits within each brain region and animal ID. If fewer than 3 images were retained, the animal ID was discarded from further analysis for the specific region. In the next step, all variables were averaged for each animal ID and region after which a similar ID-based outlier detection was applied. For longitudinal assessment of all markers, Two-way ANOVA was used with main factors of genotype and age and genotype*age interaction. In the case of interaction, post hoc comparisons (FDR corrected for 10 comparisons, p < 0.05) were performed for values within each genotype across ages compared to a control age of 3 months and between genotypes per age. When no interaction was present, the model was recalculated only for the main effects and a post hoc comparison (FDR, p < 0.05) was performed for each effect separately (for age, comparisons are made to 3 months as a control). In the case of 2B4 occupancy in CD13+ and GFAP+ cells in the zQ175DN HET, two-way ANOVA included main factor age and region and their interaction, where posthoc comparisons (FDR corrected for 28 comparisons, p < 0.05) were performed for each region pair per age. All statistical analyses and graph visualizations were performed using GraphPad Prism (version 9.4.1 for Windows, GraphPad Software, San Diego, California USA, www.graphpad.com). Outlier detection was performed in R.

## RESULTS

### Representative short (LCN and DMLN) and long (LCN DMLN) QPPs

Spatiotemporal hierarchical clustering of short (3s) QPPs revealed two robust clusters in both WT and zQ175 HET mice at every age (Fig. S1A-B). As previously reported, short QPPs represent distinct singular brain states in the rodent brain^18,22,23,31^. The QPPs of the first cluster showed co-activation of regions that represent the LCN, such as MCtx 1, MCtx 2, S1Ctx^b^, S2Ctx, lCPu, frontal association cortex (FrACtx) and insular cortex (InsCtx) and a simultaneous co-deactivation of regions that represent the DMLN like CgCtx, RspCtx, pre-infralimbic cortex (PrL/ILCtx), mCPu, S1Ctx^a^, dentate gyrus (DG), entorhinal cortex (EntCtx), auditory cortex (AuCtx) and olfactory area (OA). The QPPs belonging to the second cluster exhibited inverse activity with the co-activation of DMLN and co-deactivation of LCN regions. These short QPPs are further named LCN and DMLN QPP, based on the co-activated network they display (Fig.1A). To understand a more complex brain state periodicity, long (10s) QPPs were extracted. Clustering revealed long QPPs that constitute an interchange of activity between the two dominant networks, where LCN transitions into DMLN activation (Fig.1B). This QPP is the rodent homolog of the human primary QPP^19^. Hence, characteristic short and long QPPs are present in both WT and zQ175DN HET mice.

**Figure 1.**
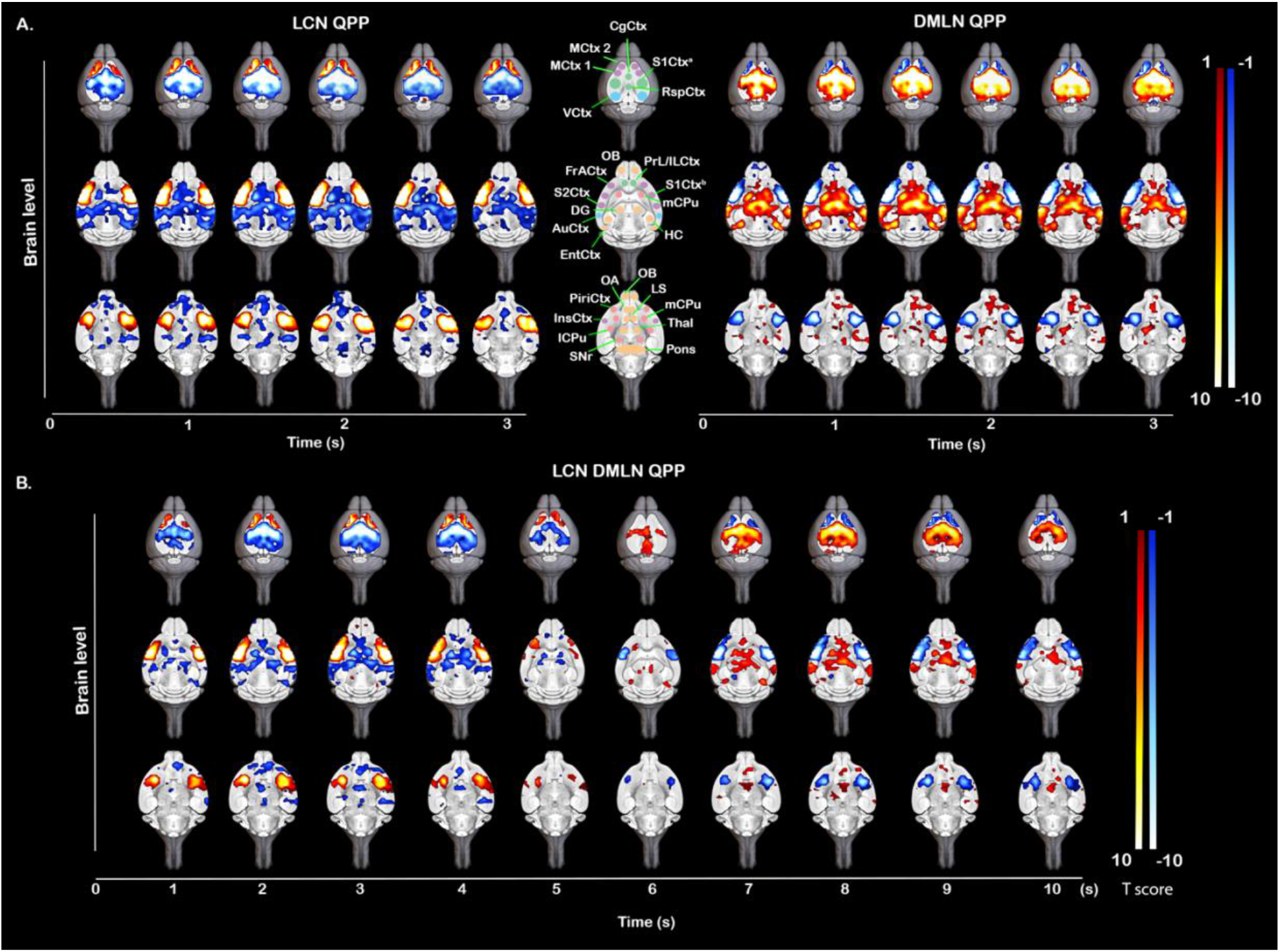
Anatomical representation of (de)activations in LCN, DMLN and LCN DMLN rQPPs. Visualization of selected brain levels of the group-wise One-sample T-maps (two-tailed one-sample T-test, FDR corrected, p < 0.05, k ≥ 10) of (A) short (3s) representative (r) LCN and rDMLN QPP (left and right) and (B) long (10s) rLCN DMLN QPP that represent significantly higher (red-yellow)/lower (blue-white) mean global BOLD activity. Maps are overlaid on a high-resolution MRI anatomical 3D image. Anatomical outline of regions corresponding to (de)activated areas in the rQPPs (A, middle). where regions are color coded whether they pertain to a specific network (green – DMLN, purple - LCN, blue - ACN, pink - SuCN) or not (orange); CgCtx cingulate cortex, MCtx 1 motor cortex 1, MCtx 2 motor cortex 2, S1Ctx^a^ somatosensory cortex 1 (trunk and lower limb), S1Ctx^b^ somatosensory cortex 1 (mouth), RspCtx retrosplenial cortex, VCtx visual cortex, OB olfactory bulb, PrL/ILCtx pre-/infralimbic cortex, FrACtx frontal association cortex, DG dentate gyrus, HC hippocampus, S2Ctx somatosensory cortex 2, mCPu – medial caudate putamen, lCPu lateral caudate putamen, AuCtx – auditory cortex, EntCtx entorhinal cortex, OA olfactory area, LS lateral septal nucleus, PiriCtx piriform cortex, InsCtx insular cortex, Thal thalamus, SNr substantia nigra reticular part;

### Age-specific spatial and temporal alterations in short (3s) LCN and DMLN QPPs

Voxel-wise differences were assessed based on the spatial properties of rQPPs, where two types of between-group comparisons were conducted for the LCN and DMLN QPPs: first, differences in the spatial (de)activation between rQPPs from each group and the second, differences in the spatial (de)activation between each rQPP of one group and its projection in the other group. The first comparison accounted for differences in most prominent QPPs that were deemed to be similar based on visual inspection and spatial correlation while the second accounted for changes in (de)activation of group representations of the same QPP (native rQPP of one group vs. its projected pattern in the other group).

At 3 months of age, a comparison between the LCN rQPPs (Fig.2A) revealed reduced activation of the ventral portion of the caudate CPu (vcCPu) and VCtx while a higher activation was present in the S1Ctx, lCPu, InsCtx and PiriCtx in the zQ175DN HET mice as compared to WT mice (Fig. 2B-left). Higher deactivation in the zQ15DN HET was found in the MCtx 2, CgCtx and the RspCtx and a portion of the VCtx (Fig.2B-left). Comparing WT LCN rQPP with its projection in zQ175DN HET, similar differences were observed (Fig.2B-right). On the other hand, the zQ175DN HET LCN rQPP revealed mainly decreased activation and deactivation in vcCPu and CgCtx respectively, when compared with its projection in WT (Fig.S2-left). The group-specific DMLN rQPP (Fig. 2C) revealed reduced activation in the RspCtx, CgCtx and VCtx and reduced deactivation in the MCtx 1, vcCPu, olfactory bulb (OB) and an inferior portion of InsCtx in the zQ175DN HET as compared to WT (Fig. 2D-left). Interestingly, an increased deactivation was present in S1Ctx, S2Ctx, lCPu and superior portions of InsCtx in the zQ175DN HET group, the same regions that showed increased activation during the LCN rQPP, indicating a potential hyper-activity compensatory mechanism. Comparison of the WT DMLN rQPP between genotypes revealed fewer differences with only decreased activation in the inferior portion of InsCtx and higher and lower deactivation in S1Ctx and MCtx 2, respectively (Fig. 2D-right). zQ175DN HET DMLN rQPP differences were only restricted to reduced activation in VCtx (Fig. S2-right).

**Figure 2.**
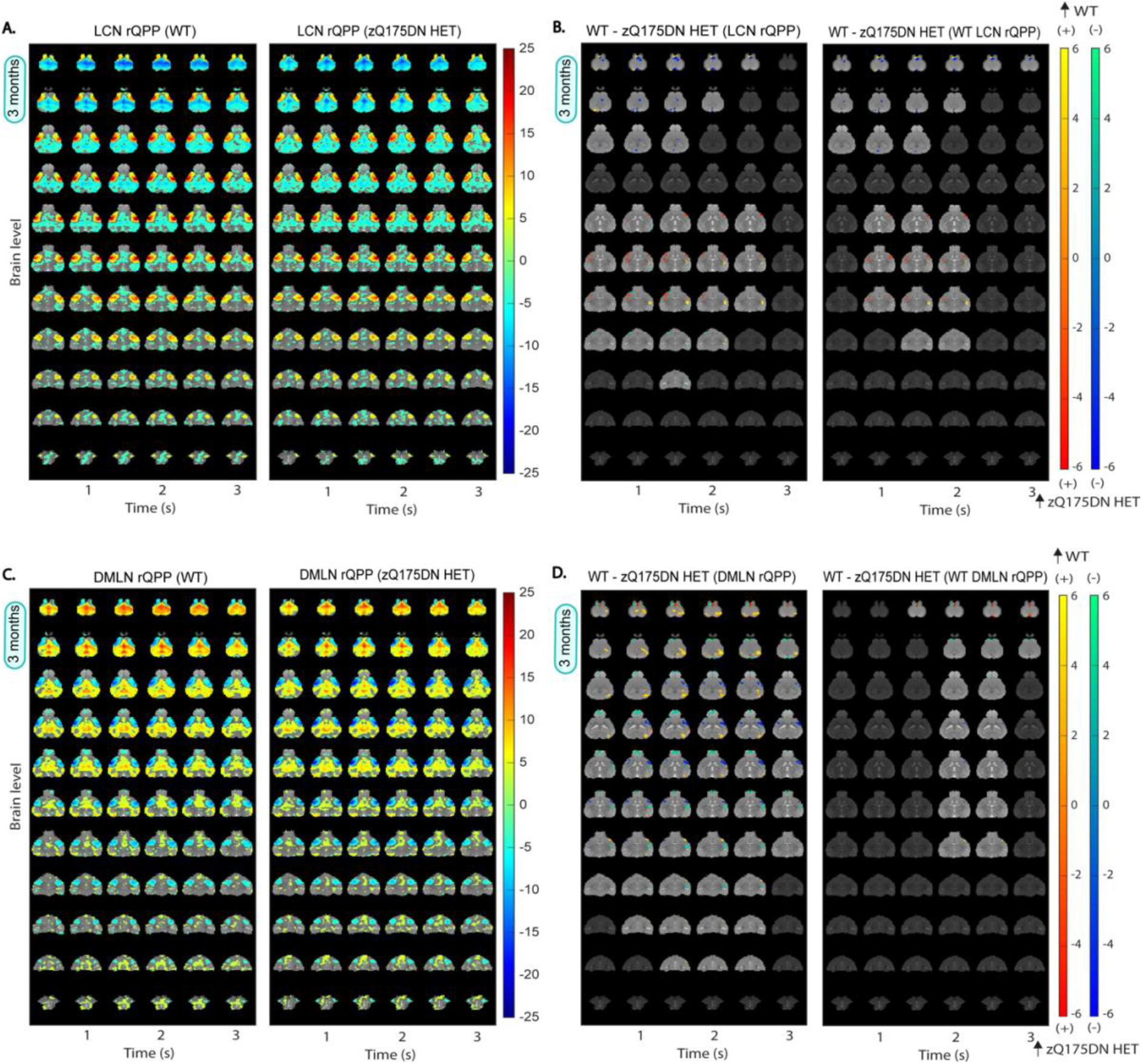
Group DMLN and LCN rQPPs in WT and zQ175DN HET and spatial activation differences at 3 months of age. A, C: One-sample T-maps of LCN (A) and DMLN (C) rQPPs in both groups where significantly activated (red-yellow) and deactivated (green-blue) voxels as compared to the mean BOLD signal are shown (two-tailed one-sample T-test, FDR, p < 0.05) B, D: Two-sample T-test maps (two-tailed, FDR, p < 0.05) of (1) between-group differences for LCN and DMLN rQPP of each group (left panels) and (2) differences between WT LCN and WT DMLN rQPPs and their projections in the zQ175DN HET (right panels). Red-yellow & blue-green colour bars display differences in voxels that are positively and negatively activated, respectively.

At 6 months of age, differences in group-based LCN rQPP (Fig. S3A) revealed increased activation in the lCPu, S1Ctx, InsCtx and PiriCtx in the zQ175DN HET (Fig. 3A-left), as at 3 months of age (Fig. 2B), and reduced deactivation in RspCtx. Moreover, a robust reduced activation was observed bilaterally in the S2Ctx and some portions of S1Ctx, also found when WT and zQ175DN HET-specific LCN rQPPs were compared with their projections in the other group (Fig. 3A-right; Fig. S3B). The group-based DMLN rQPP (Fig. S3C) comparison showed alterations in zQ175DN HET with an overall decreased activation in DMLN-related regions (RspCtx, CgCtx, hippocampus (HC)), VCtx and OB, while an increased activation was observed in the thalamus (Thal), mCPu, nucleus accumbens (NAcc) and in the superior portion of the superior colliculus (SC; Fig. 3B-left). Additionally, reduced deactivation was found in the lCPu, S1Ctx and lower portion of SC. DMLN-region diminished activation was also present in the WT or zQ175DN HET-based DMLN rQPP comparisons with their projections in the other group (Fig. 3B-right; Fig. S3D), however, the remaining differences as observed in the group-based rQPP, here, were less present.

**Figure 3.**
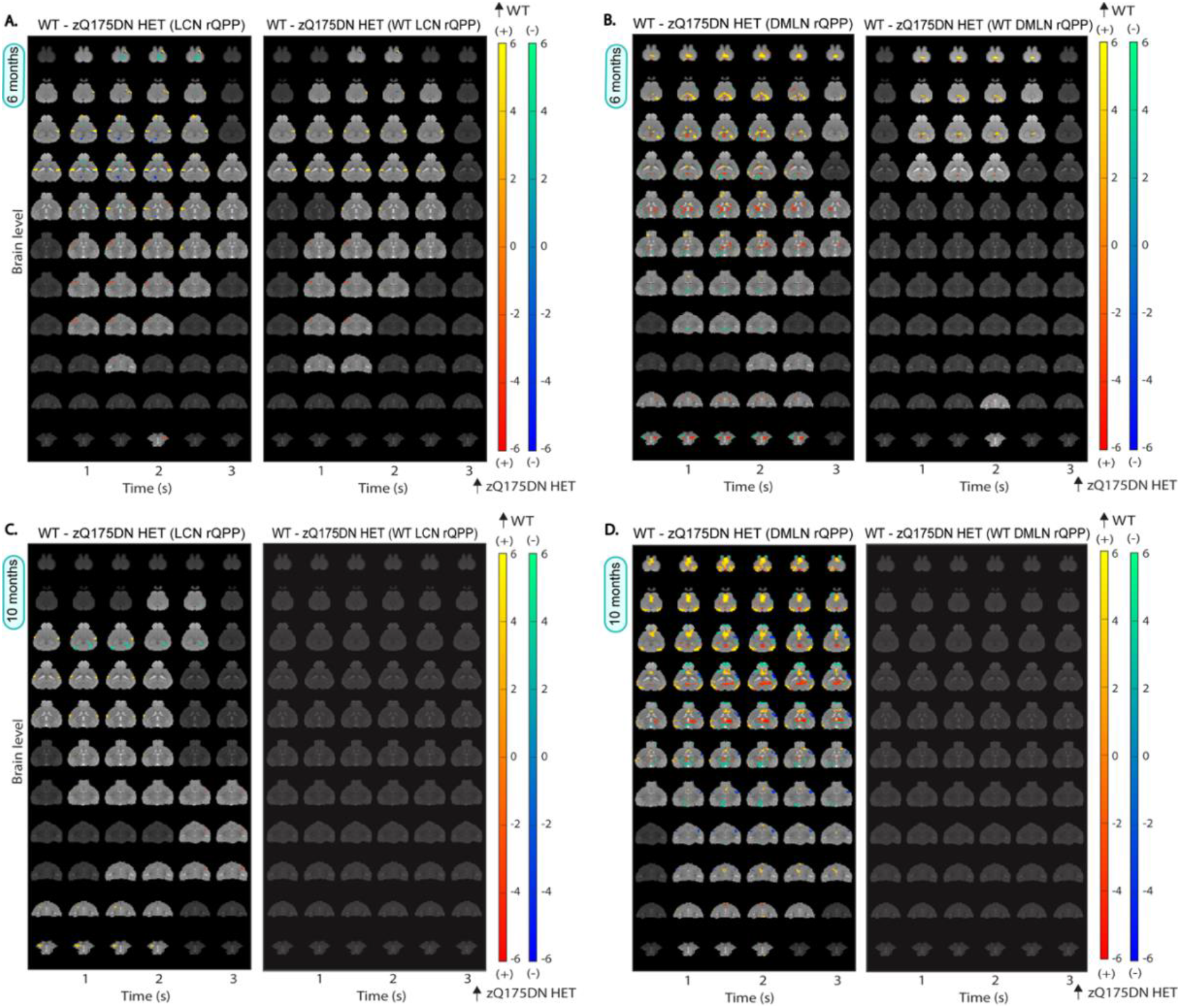
Spatial activation differences between WT and zQ175DN HET in LCN and DMLN QPP at 6 and 10 months of age. Two-sample T-test maps (two-tailed, FDR, p < 0.05) of (1) between-group differences for LCN and DMLN rQPP of each group (left panels) and (2) differences between WT LCN and WT DMLN rQPPs and their projections in the zQ175DN HET (right panels) at 6 (A, B) and 10 (C, D) months of age. Red-yellow & blue-green colour bars display differences in voxels that are positively and negatively activated, respectively.

At 10 months of age, along with the reduced activation observed bilaterally in the S2Ctx, as it was at 6 months, a reduction in activation in NAcc was also found (Fig. 3C-left) in the zQ175 DN HET when LCN rQPPs from each group (Fig. S4A) were directly compared. Higher activation was found in the lCPu and reduced deactivation in the VCtx and HC. Similar changes were found when zQ175DN HET-based rLCN QPP was compared with its projection in WT (Fig. S4B), but WT LCN rQPP showed no significant differences when compared with its projection in zQ175DN HET (Fig. 3C-right). Interestingly, direct comparison of group-based DMLN rQPPs showed significant differences across the pattern, with a robustly decreased DMLN-region activation, increased activation in the Thal, increased deactivation in the S1Ctx and decreased deactivation in the OB and SC (Fig. 3D-left). These whole-brain differences indicate that this rQPP of the zQ175DN HET is a phenotypically distinct rQPP. While the WT DMLN rQPP showed no difference with its projection in zQ175 DN HET (Fig. 3D-right), the zQ175DN HET DMLN rQPP showed several differences with reduced activation in CgCtx and VCtx and increased activation in one portion of the Thal (Fig. S4D).

Next, we compared the temporal properties, namely the occurrences of rQPPs. Here, the comparison was made between the occurrences of rQPP from each group with that of its projection in the other. The occurrence rate of the WT LCN rQPP was significantly reduced in zQ175DN HET at 3 (p = 0.0194) and 10 (p = 0.0097; Fig. 4A, 4C) but not at 6 months of age (p = 0.2116; Fig. 4B). However, WT DMLN rQPP showed significantly reduced occurrences in zQ175DN HET at all ages (3 months: p = 0.0311; 6 months: p = 0.0012; 10 months: p = 2E-06; Fig. 4). Conversely, occurrence rates of both QPPs identified in the zQ175DN HET did not differ from those of their projections in WT at any age. This indicates that the normative WT LCN and DMLN rQPP are present and progressively diminish in occurrence in the zQ175DN HET group, while the zQ175DN HET rQPPs are present in the WT group and do occur as frequently as in the zQ175DN HET group. We conclude that genotypic differences in regional activation of LCN and DMLN rQPPs are age-related, whereas the consistently decreased occurrence of the WT rQPP indicates that the predominant network states in the zQ175DN HET group are not normative.

**Figure 4.**
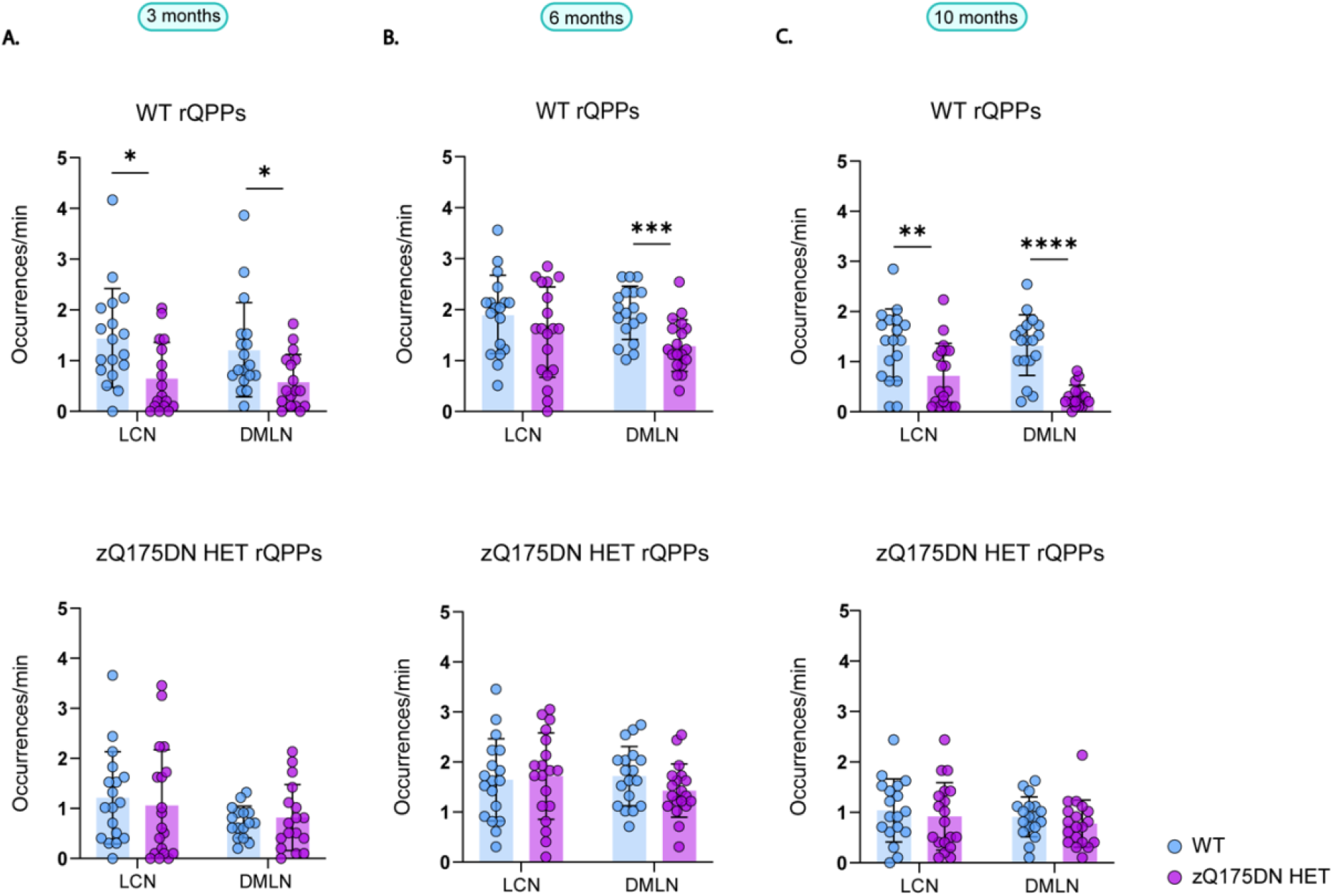
Occurrences of group-specific LCN and DMLN rQPPs within WT and zQ175DN HET at 3, 6 and 10 months of age. Between-group differences of occurrences of group-specific (WT or zQ175DN HET) rQPP with its projection in the other group (mean ± SD, two-sample T-test, FDR, p < 0.05); * p ≤ 0.05, ** p ≤ 0.01, *** p ≤ 0.001, **** p ≤ 0.0001

### LCN and DMLN rQPPs and their significance in FC alterations

As QPPs have been demonstrated as an important contributor to FC^19,22,44–47^, we investigated if the regression of the most prominent group-specific LCN and DMLN QPPs from each group will influence FC genotypic differences. At 3 months, even though no significant FC differences were observed between groups (Fig. S5A), regression of both short QPPs introduced a significantly higher FC between VCtx-S2Ctx as compared to WT (Fig. S5B). At 6 months, several pairs of regions that mostly pertain to DMLN and ACN showed a significantly reduced FC (Fig. S5C), whereas after removing the respective rLCN and rDMLN QPP, these significant FC differences disappeared, implying that these QPPs contribute to the inter-genotypic difference in FC (Fig. S5D). From the two pairs of connections that showed differences in FC at 10 months (Fig. S5E), after regression of both QPPs only the VCtx-AuCtx connection disappeared. However, several ROI pairs pertaining mostly to DMLN and ACN were now found to be significantly different between genotypes (Fig. S5F). Since the occurrence rates of both rQPPs were significantly lower in the zQ175DN HET group, regression of these QPPs had a larger impact on FCs in the WT group and resulted in higher net remnant FC in the zQ175DN HET group as compared to WT for several connections as observed in Fig. S5F. This implies that the dominant group-specific QPPs are relevant contributors to FC, as observed with the loss of genotypic differences linked to the presence of QPPs.

### Progressive spatiotemporal alterations in long LCN DMLN (and DMLN LCN) QPPs

At 3 months, hierarchical clustering of long (10s) QPPs, revealed one cluster in both groups, whose rQPP demonstrated LCN activation-DMLN deactivation pattern transitioning into DMLN activation-LCN deactivation pattern (LCN DMLN rQPP; Fig. 5A). Comparisons of voxel-wise activations of each group’s LCN DMLN rQPP were not performed due to the need for a phase shift of one group’s rQPP to be aligned to the rQPP of the other group. Since this results in a shift (typically of 1TR), the phase-aligned QPPs will not represent the true original QPP of that group. Therefore, we compared WT- and zQ175DN HET rQPPs with their projections in the other group. WT LCN DMLN rQPP showed, in the first half (LCN activation fragment), increased activation of the lCPu, decreased activation of MCtx2 and decreased deactivation of DG, in the zQ175DN HET group (Fig. 5B). However, for the zQ175DN HET LCN DMLN rQPP, the activation differences during LCN comprised of decreased activations in the MCtx2, iCPu, S1Ctx, S2Ctx and reduced deactivation in the DG as well as in the MCtx2 during DMLN activation (Fig. S7A).

**Figure 5.**
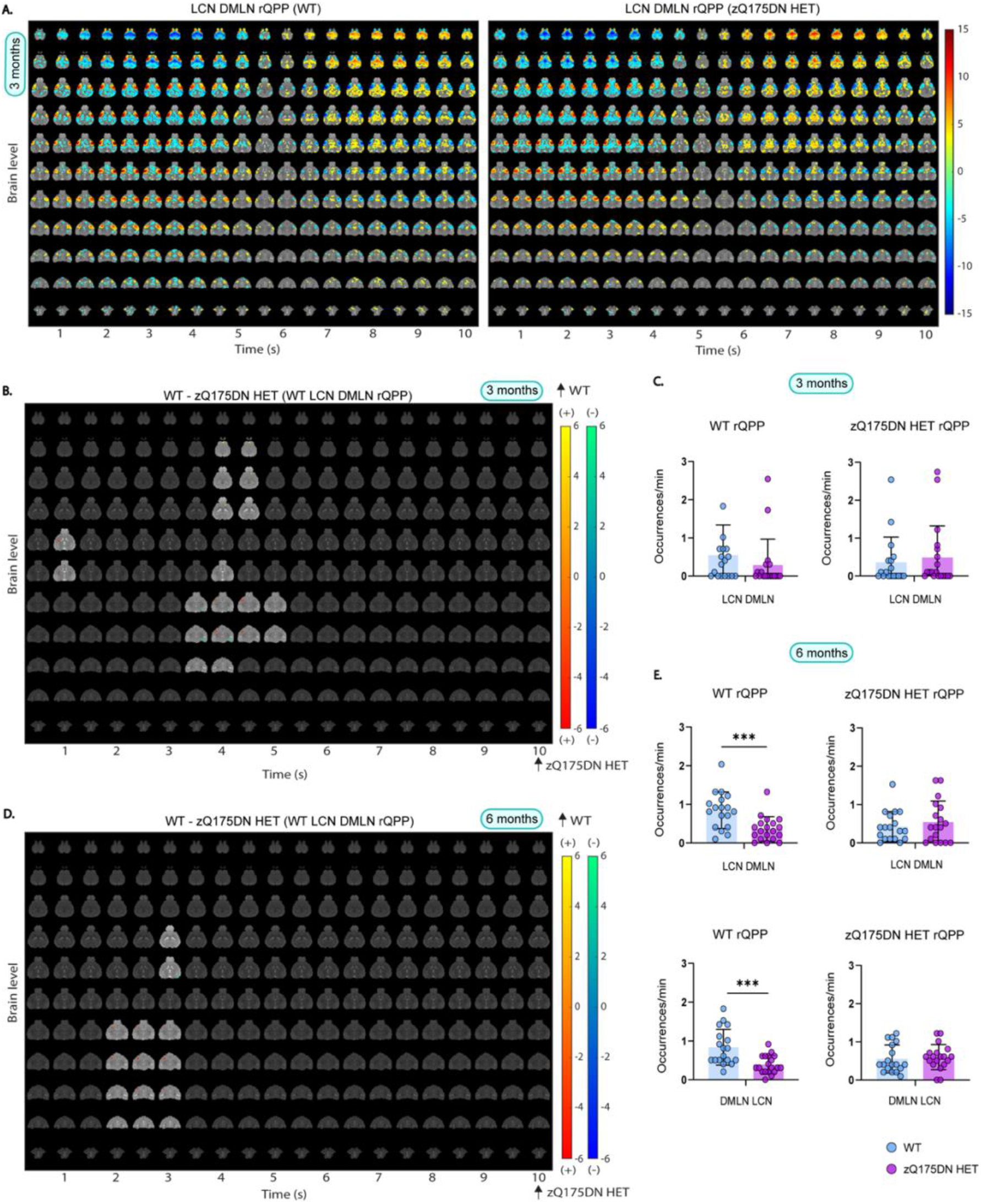
Group LCN DMLN rQPP and spatial and temporal differences in LCN DMLN and DMLN LCN QPPs at 3 and 6 months of age. (A) One-sample T-maps of rLCN DMLN QPP in both groups where significantly activated (red-yellow) and deactivated (green-blue) voxels as compared to the mean BOLD signal are shown (two-tailed one-sample T-test, FDR, p < 0.05); (B, D) Two-sample T-test maps (two-tailed, FDR, p < 0.05) of between-group differences shown for WT LCN DMLN rQPP at 3 and 6 months; Red-yellow & blue-green colour bars display differences in voxels that are positively and negatively activated, respectively. (C, E) Genotypic differences in occurrences rate for WT and zQ175DN HET LCN DMLN rQPP at 3 and 6 and DMLN LCN rQPP at 6 months (mean ± SD, two-sample T-test, FDR, p < 0.05); * p ≤ 0.05, ** p ≤ 0.01, *** p ≤ 0.001

At 6 months, two clusters were present in both genotypes, with one of them showing the LCN DMLN QPP (Fig. S6A) while the other showed an anticorrelated pattern (DMLN LCN QPP; Fig. S6B). WT LCN DMLN rQPP showed differences only during the LCN fragment, namely, increased activation in the mCPu, InsCtx and decreased deactivation in the DG. Similar differences were observed in the case of zQ175DN HET LCN DMLN rQPP with additional bilateral reduced activation of S2Ctx and iCPu (Fig. S7B). WT DMLN LCN rQPP was not different from its projection in the zQ175DN HET group, but zQ175DN HET rQPP showed, during the DMLN fragment, decreased activation of DMLN regions (RspCtx, CgCtx, HC) and the OB (Fig. S7C). At 10 months, in the WT group, one LCN DMLN cluster was present, however, in the zQ175DN HET group there was no robust cluster present but rather multiple heterogeneous clusters (Fig. S1D), suggesting that in this group, the long QPP comprising of a transition between the two anticorrelated RSNs (LCN+/DMLN-to LCN-/DMLN+) within the window of 10s is no longer present at this age.

A comparison of occurrences for the LCN DMLN rQPPs at 3 months showed no significant differences for either WT- or zQ175DN HET-based rQPP (Fig. 5C). At 6 months, WT LCN DMLN (p = 0.001) and DMLN LCN rQPPs (p = 0.001), showed a significant decrease in occurrence rates when compared with their projections, but no differences were found in the occurrence rates of zQ175DN HET rQPPs and their respective projections in WT (Fig. 5E). Thus, long QPPs, the homolog of the human QPP1, show age-related spatial activation differences and decreased normative occurrence of the WT rQPPs in the zQ175DN HET group. Moreover, the age-related decrease of the DMLN leads to a breakdown of the LCN DMLN flux state in the zQ175DN HET group.

### Regional timing differences during long LCN DMLN (and DMLN LCN) QPPs

To understand the possible driving mechanisms of coordinated activity in brain dynamics represented by the long QPPs, we investigated the relationship between timings of six prominent regions’ peak of activity during both LCN and DMLN fragments, similar to what has been shown in humans^19^. We hypothesized that due to the already observed alterations in the QPP properties, activity propagation would also be altered in the zQ175DN HET mice. Here, for 3 and 6 months, we compared regional peak timings and inter-regional peak timing relationships for each of the two peaks of activation in the WT long rQPPs with their projections in the zQ175DN HET. At 3 months, in the WT LCN DMLN rQPP, during the LCN fragment, the mean peak timing of mPFC (p = 0.0111), MCtx (p = 0.003) and RspCtx (p = 0.0113) showed a significant delay in zQ175DN HET as compared to WT (Fig. S8A). When assessing the inter-regional relationship in peak timing, in both groups, the MCtx succeeded in all other regions except mPFC in the zQ175DN HET (Fig. S8B). Moreover, in both groups, CPu and S1Ctx preceded the mPFC, however, only in the WT group, both regions also peaked before the VCtx. The delayed peak in mPFC, MCtx and RspCtx in the zQ175DN HET group led to significantly longer inter-regional peak intervals with the VCtx as compared to WT (Fig. S8C). During DMLN activation, MCtx (p = 7.36E-05) and S1Ctx (p = 0.0089) had a significantly delayed peak of de-activation (Fig. S8D) in zQ175 DN HET. In WT, mPFC and the VCtx preceded all other regions and CPu peaked significantly before MCtx, while the S1Ctx peaked last, but within the zQ175DN HET group, all other regions peaked before both MCtx and S1Ctx (Fig. S8E). Intergenotypic comparisons revealed that the MCtx had altered inter-regional peak relationship with all other regions in the zQ175DN HET as compared to WT (Fig. S8F).

At 6 months of age, timings were compared for WT LCN DMLN and DMLN LCN rQPPs. In the WT LCN DMLN rQPP, during the LCN activation, both S1Ctx (p = 3.04E-05) and RspCtx (p = 0.0037) showed a significantly earlier mean peak timing (Fig. 6A). Within WT, a significant difference in peak timing was observed for the CPu peak that precedes S1Ctx, while within zQ175DN HET the VCtx is preceded by the S1Ctx and the RspCtx which also peaks before mPFC and the MCtx (Fig. 6B). Genotypic differences showed that the premature peak timings in S1Ctx and RspCtx led to altered inter-peak intervals with all other regions in the zQ175DN HET (Fig. 6C).

**Figure 6.**
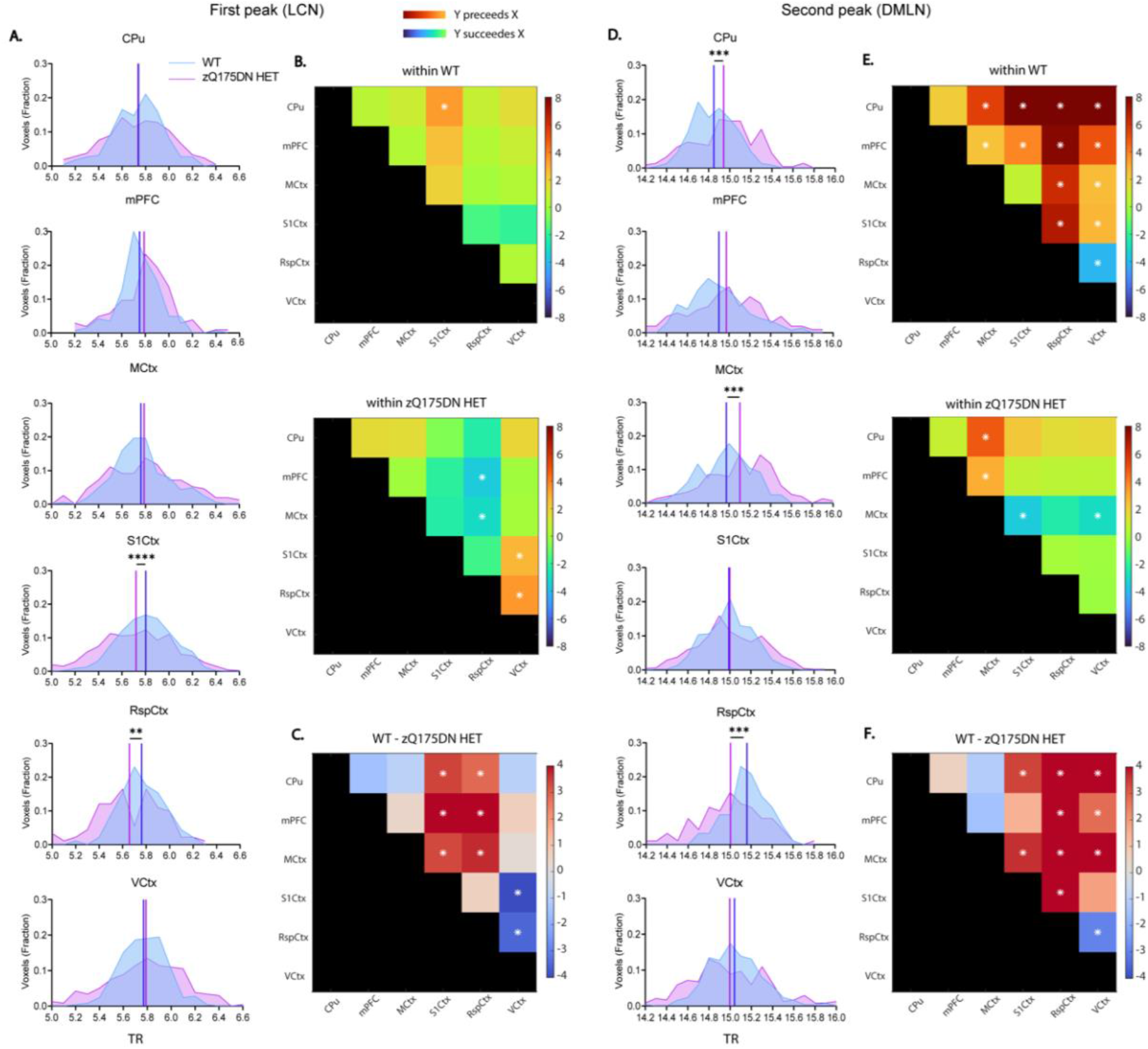
Regional mean timing and inter-regional timing differences in WT LCN DMLN rQPP at 6 months of age. (A, D) histogram of the distribution of significant voxels with the between-group differences in mean peak timings for each region during the first (LCN) and second (DMLN) peak of WT LCN DMLN rQPP; (B, E) between region timing relationship within each group; (C, F) between-group differences of inter-regional timing intervals; All comparisons were corrected for both metrics and all regions (two-sample t-test, FDR, p < 0.05); * p ≤ 0.05, ** p ≤ 0.01, *** p ≤ 0.001, **** p ≤ 0.0001; CPu caudate putamen, mPFC medial prefrontal cortex, MCtx motor cortex, SS1Ctx Somatosensory Cortex 1, RspCtx retrosplenial cortex, VCtx visual cortex;

During DMLN activation, both CPu (p = 3.72E-04) and the MCtx (p = 3.0E-04) showed a significantly delayed peak of deactivation while the RspCtx (p = 3.0E-04) had a significantly earlier peak timing in the zQ175DN HET (Fig. 6D). Within WT, CPu and mPFC preceded the peak timings of all other regions, while the VCtx and RspCtx peaked after all other regions with RspCtx peaking also after the VCtx (Fig. 6E-top). However, within the zQ175DN HET group, the MCtx has a significantly lagged peak after all other regions except RspCtx (Fig. 6E-bottom). Between-group comparisons showed an altered relationship of the RspCtx with all other regions, but also of the VCtx with other regions except S1Ctx which in turn showed a significantly different peak timing relationship with CPu and MCtx in the zQ175DN HET as compared to WT (Fig. 6F).

In the case of WT DMLN LCN rQPP, where the DMLN was first activated, in the zQ175DN HET, a significantly delayed S1Ctx peak of deactivation (p = 2.03E-04) with an earlier peak of RspCtx (p = 0.0394) activation was observed (Fig. S9A). In both groups, the VCtx peaked last. Additionally, in the zQ175DN HET, the S1Ctx succeeded all other regions (Fig. S9B). Moreover, the CPu had an earlier peak compared to mPFC in the WT group. The lagged peak of the S1Ctx revealed altered inter-regional peak timing differences with mPFC, the VCtx and the RspCtx in zQ175DN HET with respect to the WT (Fig. S9C). During LCN activation, a significantly delayed peak of CPu (p = 0.0394) and an earlier RspCtx peak (p = 0.0394) were present in the zQ175DN HET group (Fig. S9D). In both groups, the mean peak timing of CPu preceded all other regions except mPFC in the zQ175DN HET (Fig. S9E). Moreover, mPFC and S1Ctx peaked before the MCtx and the RspCtx peak lagged all other regions in both groups. Genotypic differences were observed in the altered relationship of CPu with RspCtx in the zQ175DN HET group (Fig. S9F) due to the inverse directionality of inter-genotypic change in the peak timings of these two regions. These results reveal that the nuanced timing relationship of different regions is affected in the zQ175DN HET group relative to WT. Specifically, a premature peak of the RspCtx regardless of the network state, indicates asynchrony of this region with respect to the others involved in the long QPPs.

### mHTT deposition in pericytes across different ages

To find out whether the progressive deposition of mHTT correlated with the observed defects in QPPs, we quantified the 2B4-positive spot coverage per cell (Fig. 7A). The 2B4 spot coverage showed a significant genotype x age interaction (Two-way ANOVA, *p* < 0.05) for all considered brain regions. Post-hoc comparisons (FDR, *p* < 0.05) showed an increase of 2B4 spot coverage from 6 months onwards in both mCPu and lCPu but also in the InsCtx, PiriCtx and MCtx 2, while a significant increase in 2B4 spot coverage from 8 months onwards was found in MCtx 1, S1Ctx and CgCtx in the zQ175DN HET group (see table S1; Fig. 7B). When refining the analysis to pericytes, by limiting the quantification of the 2B4 spot coverage to CD13-positive cells, we found no significant difference at 3 months, but a significant increase at 6 months in mCPu, S1Ctx and MCtx 2 (Fig. 7D). At 8 and 12 months of age, only mCPu and lCPu showed a significantly increased 2B4 spot coverage of CD13-positive cells as compared to the other regions (Fig. 7D). This suggests that the CPu shows consistently increase in mHTT load across ages in pericytes, however, cortical regions show only a transient increase in mHTT load. Of note, these changes were independent of pericyte abundance since no significant differences were observed in the total CD13 area for any region (Fig. S11A). Genotypic comparison of the number of CD13-positive cells revealed an interaction in lCPu and S1Ctx where post-hoc comparisons showed that only at 6 months of age there was a trend of decrease (*p* = 0.063) and a significant decrease (*p* = 0.035) in cell count in the zQ175DN HET, respectively (Fig. 7C). In the MCtx 2, an overall higher cell count across ages was observed in zQ175DN HET as compared to WT. Together, these results demonstrate that, despite the lack of difference in total pericyte area, cortical regions show changes in pericyte abundance.

**Figure 7.**
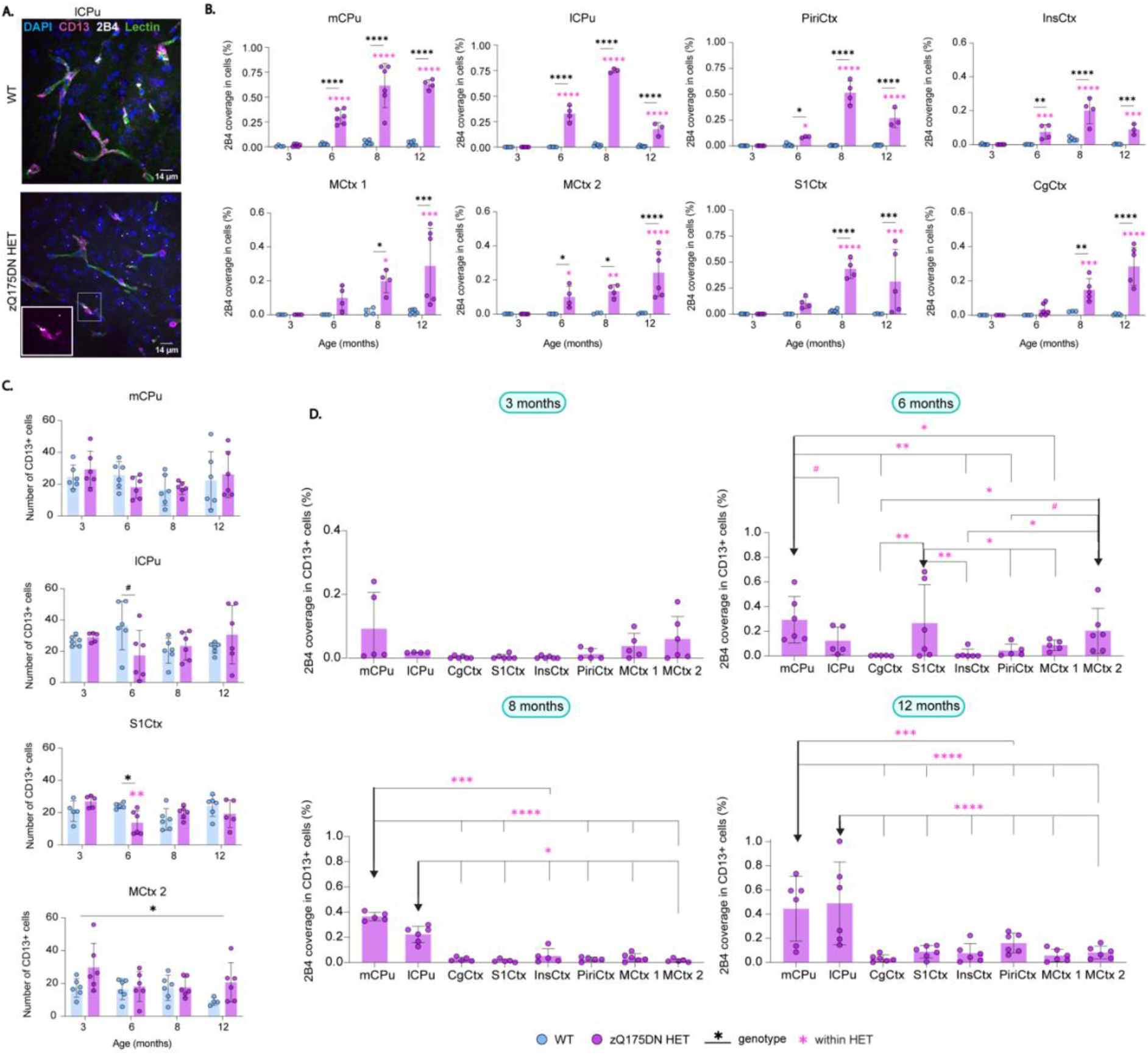
mHTT deposition in total cells and pericytes in zQ175DN mice across ages. (A) Immunofluorescence representative image for mHTT (2B4, white), pericytes (CD13, magenta) aligning blood vessels (lectin, green) and nuclei (DAPI, blue) in WT and zQ175DN HET in the lCPu at 6 months of age; between-group comparison of (B) 2B4 coverage in all cells based on 2B4 spot total area relative to the cell area and (C) number of CD13+ cells across ages (mean ± SD, two-way ANOVA, FDR, p < 0.05); (D) deposition of 2B4 in CD13+ cells in zQ175DN HET mice across regions at 3, 6, 8 and 12 months of age (mean ± SD, two-way ANOVA, *p* < 0.05); post-hoc comparisons represented as differences to a specific region (**↓**), FDR corrected, \**p ≤* 0.05, ** *p* ≤ 0.01, *** *p* ≤ 0.001, **** *p* ≤ 0.0001, # *p* ≤ 0.1; mCPu medial caudate putamen, lCPu lateral caudate putamen, PiriCtx piriform cortex, InsCtx insular cortex, MCtx 1 motor cortex 1, MCtx 2 motor cortex 2, S1Ctx somatosensory cortex 1, CgCtx cingulate cortex;

### mHTT deposition in astrocytes and VEGF changes across different ages

To assess the mHTT occupancy within astrocytes, changes in 2B4 spot coverage within GFAP+ cells in the zQ175DN HET mice between regions per age were evaluated (two-way ANOVA, *p* < 0.05). Whereas no differences were present at 3 months, astrocyte 2B4 spot coverage at 6 months, revealed that the mCPu was the only region that showed a significant increase in the 2B4 area occupied within GFAP cells, as compared to CgCtx and MCtx 2 (Fig. 8B). At 8 and 12 months of age, both the mCPu and PiriCtx had a significantly increased 2B4 spot coverage in GFAP+ cells as compared to the other regions (Fig. 8B). Additionally, as a measure of reactivity in astrocytes, we calculated the GFAP area (Fig. 8C), where only the lCPu showed an interaction with a significant increase in zQ175DN HET at 12 months (p = 0.01), whereas an overall increase in the InsCtx was present across ages. GFAP+ cell count comparison revealed no significant interaction for any of the regions, except for the InsCtx, with an overall increase in GFAP cell count in the zQ175DN HET mice (Fig. S12B). Altogether, we observe that mHTT load in astrocytes is consistently higher in the CPu and later on in the PiriCtx, as compared to other regions, whereas changes linked to astrogliosis were steadily present only in the InsCtx. VEGF has a direct association with both pericytes and astrocytes^48–50^, and its increase has been defined as a neuroprotective response to glutamate excitotoxicity, a prevailing hypothesis in HD^51–53^. Therefore, we measured the mean VEGF intensity to assess changes in this factor. From the 8 regions we investigated, lCPu, CgCtx, S1Ctx, InsCtx and PiriCtx had a significant interaction between genotype and age (see Table S8). Post-hoc comparisons (FDR, *p* < 0.05) in these regions showed a markedly significant increase at 6 months in the zQ175DN HET group as compared to WT (Fig. 8D). In addition, CgCtx showed a significant VEGF increase also at 8 months and the lCPu at 12 months of age. Overall, the pronounced increase in VEGF in the zQ175DN HET mice at 6 months could suggest a compensatory effect that coincides with the reported onset of motor deficits in this model^26^.

**Figure 8.**
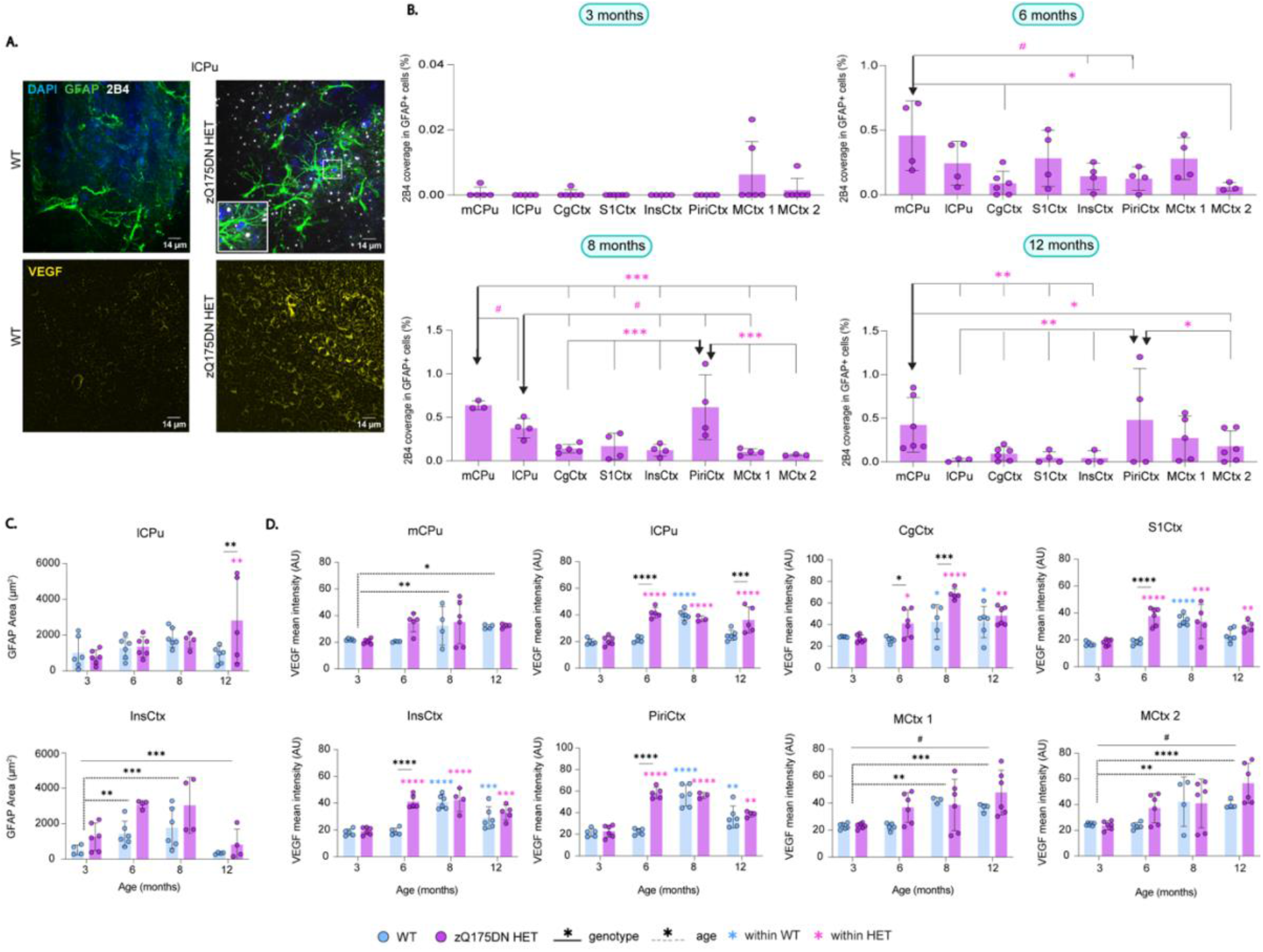
mHTT deposition in astrocytes and VEGF expression in the zQ175DN HET mice across ages. Representative immunofluorescence images of (A, top) mHTT (2B4, white), astrocytes (GFAP, green) and nuclei (DAPI, blue) at 8 months of age and (A, bottom) an angiogenesis marker (VEGF, yellow) at 6 months of age for WT and zQ175DN HET; (B) deposition of 2B4 in GFAP+ cells in zQ175DN HET mice across regions at 3, 6, 8 and 12 months of age (mean ± SD, two-way ANOVA, p < 0.05); genotypic difference in (C) thresholded GFAP+ area relative to FOV and in (D) VEGF intensity in FOV (mean ± SD, two-way ANOVA, *p* < 0.05); post-hoc comparisons represented as differences to a specific region (**↓**), FDR corrected, * *p* ≤ 0.05, ** *p* ≤ 0.01, *** *p* ≤ 0.001, **** *p* ≤ 0.0001, # *p* ≤ 0.1; AU arbitrary unit, mCPu medial caudate putamen, lCPu lateral caudate putamen, PiriCtx piriform cortex, InsCtx insular cortex, MCtx 1 motor cortex 1, MCtx 2 motor cortex 2, S1Ctx somatosensory cortex 1, CgCtx cingulate cortex;

## DISCUSSION

Our study is the first comprehensive study of the evolution of dynamic recurring whole-brain states in an HD model, represented by QPPs, which are relevant contributors to FC. These findings characterize the fine scale of instantaneous changes in the BOLD-based alterations in HD, which show the aberrant evolution of the dynamics of the most prominent resting-state networks – the DMLN and the LCN. An initial region-specific hyperactivity in these QPPs was observed as an early compensation to maintain normal brain network function, before any brain atrophy or motor disturbances are present, as reported in this HD model^26,54^. In addition, we found progressive cessation of the DMLN activity that led to a loss of the presence of the rodent homolog of the human primary QPP. These alterations occur when the motor deficits and brain atrophy are present^26,54^, suggesting that the loss of network integrity could contribute to the phenotype progression in this HD mouse model. Moreover, the reduced occurrence of the normative QPPs of healthy controls in the HD mouse model suggests that QPP projection holds translational potential as a method to inform on the different disease stages in PwHD. With this study we have described the dynamic whole brain network changes along phenotypic progression in an HD animal model, which can open new avenues of investigation when applied in clinical HD research. Furthermore, our work demonstrates that measuring dynamic brain states is a significant improvement over the current static FC methods, not only for preclinical investigation but also potentially for clinical HD research.

### Diminished DMLN activity - an instigator for a breakdown of the LCN DMLN activation cycle

Previously, we found differences in FC starting in DMLN and ACN regions in the zQ175DN HET mice as compared to WT, at 6 months of age ^29^. Here, we reveal dynamic QPP states already at 3 months with increased deactivation, and decreased activation in parts of the RspCtx, CgCtx, MCtx 2 and VCtx, all regions that are part of the core activation of DMLN^55^. Moreover, these regions showed a lagged peak of deactivation during LCN in the LCN DMN QPP, while the MCtx also lagged in its peak of activation in the DMLN segment, leading to altered relationships with the other regions. Deficits in neuronal synchrony in the motor cortex during non-motion, measured with Ca^2+^ imaging, as well as electrophysiological alterations, were also reported at this age in the zQ175 mice, explaining a potential compensatory mechanism of motor network synchrony during motion^56,57^. At 6 months of age, our findings show a prominent reduction in BOLD de- and activation of the RspCtx in LCN and DMLN QPP, respectively, accompanied by reduced activation also in CgCtx and VCtx. This marked decrease in activation in the DMLN QPP was also seen in the reduced occurrence of this pattern in the zQ175DN HET mice. Remarkably, the RspCtx also showed a premature peak of either activation or deactivation in both long QPPs, indicating that there is an asynchronous peak in the BOLD activity that leads to altered time-locked relationships with other regions. A recent study in zQ175DN mice of the same age reported altered mesoscale cortical activity in the RspCtx during a behavioral task where they report that initial behavioral deficits occur at this age, suggesting a potential role of the RspCtx in phenoconversion in this model^58^. The overall diminished DMLN activity at 10 months also led to a breakdown of the LCN DMLN state flux reflected in the lack of presence of the LCN DMLN QPP in the zQ175DN HET group, alongside the decreased occurrence of DMLN. Human FC has been shown to rely on a particular hierarchy that is reflected in cortical gradients of the RSNs, which shows variation from unimodal regions on one end, such as S1Ctx and S2Ctx, which are hard-wired and have stronger structural-functional coupling, and transmodal regions on the other end, which underly multi-input integration, such as the DMN constituents, a major hub for incorporating cortical information^59–62^. This gradient cortical hierarchy was also found in mice, with similarities observed in both functional and long-range axonal properties^61,63,64^. The transmodal DMLN regions play a central part in multimodal input, hence, they are highly adaptive to different types of information, therefore playing a crucial role in normal sensory processing and integration^62,63^. However, in pathological conditions, these regions are more vulnerable to processes of neurodegeneration; this was observed in Parkinson’s disease patients, where DMN shows higher structural-functional decoupling in DMN, which was also linked to a decrease in dopamine transmission^65^. Our observations from both FC and dynamic analysis point to the DMLN as the most vulnerable network in the zQ175DN HET that leads to a progressive prevalence of segregated states that pertain to the hard-wired unimodal regions. To be able to better link the structural coupling to these changes, future investigations of T1w:T2w MRI could help in mapping the myelin-based gradients in the cortex^63,66^.

### Non-normative QPPs govern brain-wide activity – a case for the neurodevelopmental hypothesis

We observed that the normative WT-based QPPs are ubiquitously underrepresented at all ages in the zQ175DN HET group. Surprisingly, the zQ175DN HET-based QPPs were present in the WT with the same occurrence. This suggests that the most dominant forms of DMLN and LCN QPPs in the zQ175DN HET group are a weaker state of these RSNs, as they still occur with the same frequency within the WT, but less than the normative QPPs. This implies a possible neurodevelopmental role underlying the RSNs in this model. A growing body of evidence supports the hypothesis that HD has neurodevelopmental origin^67–71^. This hypothesis posits that the mutated huntingtin gene alters specific brain regions and their circuits from birth, but these maladaptations are compensated for in early life, while these abnormal cells are in a quiescent-like state^69^. During adulthood, normal ageing and epigenetic factors influence the breakdown of this protective mechanism which leads to the known neurodegenerative process. Many HD models have demonstrated cortical maldevelopment to have a short-term change in a specific postnatal window that shows transient circuitry alterations but also brain volume abnormalities, which were observed in zQ175 male mice^72–74^. As regional volume postnatal changes are present in this model^74^, assessing how this impacts RSNs in this critical period would aid in better understating of the developmental effects on whole brain circuitry in HD^70,75^.

### mHTT deposition in astrocytes and pericytes linked to QPP alterations

The role of astrocytes and their dysfunction in HD pathology is gaining attention as their impairment has been postulated to play a critical role in imbalanced glutamate homeostasis, the underlying cause of excitotoxicity in HD^76,77^. For the first time, we have characterized GFAP+ astrocytes across different cortical regions in this model, where at 6 months, despite no change in the total area which is indicative of astrogliosis, we observed an overall increase in 2B4 spot coverage with the mCPu showing the highest percentage. This increase in 2B4 spot coverage was also observed at 8 months in mCPu, lCPu and PiriCtx. In GFAP+ astrocytes, an increase in mHTT inclusion diameter was observed at the same age in the frontal cortex and striatum in this model^78^. However, S100ß+ cells, a marker for nonreactive astrocytes, showed more inclusions even from 6 months of age and it has been reported that this marker is more sensitive for staining all astrocytes^78–80^. Increases in mHTT coverage within astrocytes in the CPu and several cortical regions with the absence of astrogliosis can indicate the presence of functional alterations that can potentially influence the BOLD-related alterations observed with QPPs. Moreover, an increase in GFAP+ area and cell count in the InsCtx was present across ages, which means that astrogliosis is present in this region. It is tempting to speculate that this susceptibility is due to the InsCtx being one of the transmodal regions in the functional gradients which is more sensitive under pathological conditions, such as the phenotype driven by mHTT expression in the zQ175DN HET model^63^. Pericytes are an integral part of the neurogliovascular unit, especially at the capillary level, where they regulate the vessel diameter in response to the energy demands of the brain^81,82^. Pericytes have not been well investigated in an HD context except for two studies—one in R6/2 mice and another using cell cultures from PwHD—that show decreased pericyte coverage in the cortex^83,84^. In our study, we characterize for the first time pericyte alterations in an HD model in several cortical and striatal regions across different ages. In the zQ175DN HET group, at 6 months of age, we observed increased 2B4 spot coverage in CD13+ pericytes in mCPu, S1Ctx and MCtx 2. Moreover, only at this age, decreased pericyte count was present in the S1Ctx and a trend of decrease in the CPu at this age. It can be inferred that loss of pericytes in these regions can be a transitory change, coinciding with the phenotypic conversion in this model, that is being compensated for after initial abnormalities occur^26,58^. Interestingly, these regions show consistent hyper(de)activations in QPPs. The MCtx 2 however, has an overall increase in pericytes across ages, which could allude to a developed maladaptation in this region. The presence of increased but potentially dysfunctional pericytes can influence neurovascular coupling, which can be linked to the weaker (de)activation observed in the MCtx in QPPs. VEGF is typically known as a marker for angiogenesis, however, there are numerous functions linked to this factor and has been implicated in the etiology and treatment of neurodegenerative diseases^52,53,85,86^. In R6/2 mice and cell cultures from PwHD, increases in VEGF are observed and linked to an increase in inflammatory-prone astrocytes that leads to decreased pericyte coverage^83^. Our findings reveal a marked increase of VEGF in lCPu, S1Ctx, InsCtx and PiriCtx at 6 months of age, regions that have shown hyperactivation in QPPs. VEGF has neuroprotective effects, one of which is reducing vulnerability against glutamate excitotoxicity by inducing expression of the GluR2 subunit of the AMPA receptors^51^.

In HD, reduction of this GluR2 subunit has been observed in postmortem brain tissue in the putamen, pointing to alterations in synaptic transmission^87^. Changes in hyperactivation in these regions in QPPs are also present earlier, at 3 months, which could suggest that initial changes on the circuitry level induce a VEGF increase at 6 months, as a compensatory neuroprotective mechanism that results from aberrant corticostriatal connectivity.

### Limitations

In this study we used only zQ175 HET male mice since they exhibit brain volume changes, resembling what is observed in PwHD^74^. Since the cortico-striato-thalamic network is comprised of sub-segregated regional circuitry^88,89^, to have a more granulated assessment of how these regions are altered in their timings during QPPs, future studies should employ a detailed whole brain parcellation to unravel these changes. In our IF study, we did not observe changes in 2B4 spot coverage in cells at 3 months, despite this being shown in this model, using different markers, such as the antibodies S829 or S830, which observe changes starting even from 1-2 months of age^78,90^. Likewise, the use of S100ß as a marker for astrocytes would be a more appropriate replacement compared to GFAP, which only represents inflammatory states of astrogliosis. Moreover, we did not include the RspCtx as a region of interest– these markers should be assessed in this region as well as it clearly showed marked QPP changes. To disentangle the cell-specific alterations in brain-wide network dynamics, future studies should apply chemo- or optogenetic modulation specific to astrocytes or pericytes in combination with rsfMRI to observe *in vivo* brain-wide changes pertaining to these cell types. A complementary behavioral assessment alongside the longitudinal rsfMRI time points would allow for a reliable link to the phenotypic evolution in this model and the deficits that develop at different ages which can potentially be linked to the observed brain network changes.

## ACKNOWLEDGMENTS

We express our immense gratitude to Annemie Van Eeteveldt and Zoë Embrechts for their help with the immunofluorescence study.

## REFERENCES

1. Ghosh R, Tabrizi SJ. Clinical Features of Huntington’s Disease. Adv Exp Med Biol. 2018;1049:1–28. doi:10.1007/978-3-319-71779-1_1

2. Bates GP, Dorsey R, Gusella JF, et al. Huntington disease. Nat Rev Dis Primers. Apr 23 2015;1:15005. doi:10.1038/nrdp.2015.5

3. A novel gene containing a trinucleotide repeat that is expanded and unstable on Huntington’s disease chromosomes. The Huntington’s Disease Collaborative Research Group. Cell. Mar 26 1993;72(6):971–83. doi:10.1016/0092-8674(93)90585-e

4. Ross CA, Aylward EH, Wild EJ, et al. Huntington disease: natural history, biomarkers and prospects for therapeutics. Nat Rev Neurol. Apr 2014;10(4):204–16. doi:10.1038/nrneurol.2014.24

5. Georgiou-Karistianis N, Scahill R, Tabrizi SJ, Squitieri F, Aylward E. Structural MRI in Huntington’s disease and recommendations for its potential use in clinical trials. Neurosci Biobehav Rev. Mar 2013;37(3):480–90. doi:10.1016/j.neubiorev.2013.01.022

6. Scahill RI, Andre R, Tabrizi SJ, Aylward EH. Structural imaging in premanifest and manifest Huntington disease. Handb Clin Neurol. 2017;144:247–261. doi:10.1016/B978-0-12-801893-4.00020-1

7. Pini L, Jacquemot C, Cagnin A, et al. Aberrant brain network connectivity in presymptomatic and manifest Huntington’s disease: A systematic review. Hum Brain Mapp. Jan 2020;41(1):256–269. doi:10.1002/hbm.24790

8. Lowe MJ, Dzemidzic M, Lurito JT, Mathews VP, Phillips MD. Correlations in low-frequency BOLD fluctuations reflect cortico-cortical connections. Neuroimage. Nov 2000;12(5):582–7. doi:10.1006/nimg.2000.0654

9. Greicius MD, Krasnow B, Reiss AL, Menon V. Functional connectivity in the resting brain: a network analysis of the default mode hypothesis. Proc Natl Acad Sci U S A. Jan 7 2003;100(1):253–8. doi:10.1073/pnas.0135058100

10. Fransson P. Spontaneous low-frequency BOLD signal fluctuations: an fMRI investigation of the resting-state default mode of brain function hypothesis. Hum Brain Mapp. Sep 2005;26(1):15–29. doi:10.1002/hbm.20113

11. Zhang HY, Wang SJ, Liu B, et al. Resting brain connectivity: changes during the progress of Alzheimer disease. Radiology. Aug 2010;256(2):598–606. doi:10.1148/radiol.10091701

12. Luo C, Song W, Chen Q, et al. Reduced functional connectivity in early-stage drug-naive Parkinson’s disease: a resting-state fMRI study. Neurobiol Aging. Feb 2014;35(2):431–41. doi:10.1016/j.neurobiolaging.2013.08.018

13. Poudel GR, Egan GF, Churchyard A, Chua P, Stout JC, Georgiou-Karistianis N. Abnormal synchrony of resting state networks in premanifest and symptomatic Huntington disease: the IMAGE-HD study. J Psychiatry Neurosci. Mar 2014;39(2):87–96. doi:10.1503/jpn.120226

14. Dumas EM, van den Bogaard SJ, Hart EP, et al. Reduced functional brain connectivity prior to and after disease onset in Huntington’s disease. Neuroimage Clin. 2013;2:377–84. doi:10.1016/j.nicl.2013.03.001

15. Wolf RC, Sambataro F, Vasic N, et al. Visual system integrity and cognition in early Huntington’s disease. Eur J Neurosci. Jul 2014;40(2):2417–26. doi:10.1111/ejn.12575

16. Calhoun VD, Miller R, Pearlson G, Adali T. The chronnectome: time-varying connectivity networks as the next frontier in fMRI data discovery. Neuron. Oct 22 2014;84(2):262–74. doi:10.1016/j.neuron.2014.10.015

17. Allen EA, Damaraju E, Plis SM, Erhardt EB, Eichele T, Calhoun VD. Tracking whole-brain connectivity dynamics in the resting state. Cereb Cortex. Mar 2014;24(3):663–76. doi:10.1093/cercor/bhs352

18. Majeed W, Magnuson M, Keilholz SD. Spatiotemporal dynamics of low frequency fluctuations in BOLD fMRI of the rat. J Magn Reson Imaging. Aug 2009;30(2):384–93. doi:10.1002/jmri.21848

19. Yousefi B, Keilholz S. Propagating patterns of intrinsic activity along macroscale gradients coordinate functional connections across the whole brain. Neuroimage. May 1 2021;231:117827. doi:10.1016/j.neuroimage.2021.117827

20. Wang K, Majeed W, Thompson G, Ying K, Zhu Y, Keilholz S. Quasi-periodic pattern of fMRI contributes to functional connectivity and explores differences between Major Depressive disorder and control. 2016:

21. Abbas A, Bassil Y, Keilholz S. Quasi-periodic patterns of brain activity in individuals with attention-deficit/hyperactivity disorder. Neuroimage Clin. 2019;21:101653. doi:10.1016/j.nicl.2019.101653

22. Belloy ME, Shah D, Abbas A, et al. Quasi-Periodic Patterns of Neural Activity improve Classification of Alzheimer’s Disease in Mice. Sci Rep. Jul 3 2018;8(1):10024. doi:10.1038/s41598-018-28237-9

23. van den Berg M, Adhikari MH, Verschuuren M, et al. Altered basal forebrain function during whole-brain network activity at pre- and early-plaque stages of Alzheimer’s disease in TgF344-AD rats. Alzheimers Res Ther. Oct 10 2022;14(1):148. doi:10.1186/s13195-022-01089-2

24. Menalled LB, Kudwa AE, Miller S, et al. Comprehensive behavioral and molecular characterization of a new knock-in mouse model of Huntington’s disease: zQ175. PLoS One. 2012;7(12):e49838. doi:10.1371/journal.pone.0049838

25. Southwell AL, Smith-Dijak A, Kay C, et al. An enhanced Q175 knock-in mouse model of Huntington disease with higher mutant huntingtin levels and accelerated disease phenotypes. Hum Mol Genet. Sep 1 2016;25(17):3654–3675. doi:10.1093/hmg/ddw212

26. Heikkinen T, Bragge T, Bhattarai N, et al. Rapid and robust patterns of spontaneous locomotor deficits in mouse models of Huntington’s disease. PLoS One. 2020;15(12):e0243052. doi:10.1371/journal.pone.0243052

27. Carty N, Berson N, Tillack K, et al. Characterization of HTT inclusion size, location, and timing in the zQ175 mouse model of Huntington’s disease: an in vivo high-content imaging study. PLoS One. 2015;10(4):e0123527. doi:10.1371/journal.pone.0123527

28. Fox JG BS, Davisson MT, Newcomer CE, Quimby FW, Smith AL, eds. The Mouse in Biomedical Research*, 2nd Edition*. vol 3. Elsevier 2007:637–72.

29. Vasilkovska T, Adhikari M, Van Audekerke J, et al. Resting-state fMRI reveals longitudinal alterations in brain network connectivity in the zQ175DN mouse model of Huntington’s disease. Neurobiol Dis. Mar 22 2023:106095. doi:10.1016/j.nbd.2023.106095

30. Majeed W, Magnuson M, Hasenkamp W, et al. Spatiotemporal dynamics of low frequency BOLD fluctuations in rats and humans. Neuroimage. Jan 15 2011;54(2):1140–50. doi:10.1016/j.neuroimage.2010.08.030

31. Belloy ME, Naeyaert M, Abbas A, et al. Dynamic resting state fMRI analysis in mice reveals a set of Quasi-Periodic Patterns and illustrates their relationship with the global signal. Neuroimage. Oct 15 2018;180(Pt B):463–484. doi:10.1016/j.neuroimage.2018.01.075

32. Praet J, Manyakov NV, Muchene L, et al. Diffusion kurtosis imaging allows the early detection and longitudinal follow-up of amyloid-beta-induced pathology. Alzheimers Res Ther. Jan 9 2018;10(1):1. doi:10.1186/s13195-017-0329-8

33. Liska A, Galbusera A, Schwarz AJ, Gozzi A. Functional connectivity hubs of the mouse brain. Neuroimage. Jul 15 2015;115:281–91. doi:10.1016/j.neuroimage.2015.04.033

34. Bukhari Q, Schroeter A, Cole DM, Rudin M. Resting State fMRI in Mice Reveals Anesthesia Specific Signatures of Brain Functional Networks and Their Interactions. Front Neural Circuits. 2017;11:5. doi:10.3389/fncir.2017.00005

35. Janke AL, Ullmann JF. Robust methods to create ex vivo minimum deformation atlases for brain mapping. Methods. Feb 2015;73:18–26. doi:10.1016/j.ymeth.2015.01.005

36. Schindelin J, Arganda-Carreras I, Frise E, et al. Fiji: an open-source platform for biological-image analysis. Nat Methods. Jun 28 2012;9(7):676–82. doi:10.1038/nmeth.2019

37. De Vos WH, Van Neste L, Dieriks B, Joss GH, Van Oostveldt P. High content image cytometry in the context of subnuclear organization. Cytometry A. Jan 2010;77(1):64–75. doi:10.1002/cyto.a.20807

38. Verschuuren M, Verstraelen P, Garcia-Diaz Barriga G, et al. High-throughput microscopy exposes a pharmacological window in which dual leucine zipper kinase inhibition preserves neuronal network connectivity. Acta Neuropathol Commun. Jun 4 2019;7(1):93. doi:10.1186/s40478-019-0741-3

39. Verstraelen P, Garcia-Diaz Barriga G, Verschuuren M, et al. Systematic Quantification of Synapses in Primary Neuronal Culture. iScience. Sep 25 2020;23(9):101542. doi:10.1016/j.isci.2020.101542

40. Schmidt U, Weigert M, Broaddus C, Myers G. Cell Detection with Star-Convex Polygons. Springer International Publishing; 2018:265–273.

41. Longair MP, S.; Schindelin, J. “Tubeness” plugin for ImageJ.

42. Sato Y, Nakajima S, Shiraga N, et al. Three-dimensional multi-scale line filter for segmentation and visualization of curvilinear structures in medical images. Med Image Anal. Jun 1998;2(2):143–68. doi:10.1016/s1361-8415(98)80009-1

43. R Core Team. R: A language and environment for statistical computing. R Foundation for Statistical Computing, Vienna, Austria. 2014;

44. Thompson GJ, Pan WJ, Billings JC, et al. Phase-amplitude coupling and infraslow (<1 Hz) frequencies in the rat brain: relationship to resting state fMRI. Front Integr Neurosci. 2014;8:41. doi:10.3389/fnint.2014.00041

45. Keilholz SD, Magnuson ME, Pan WJ, Willis M, Thompson GJ. Dynamic properties of functional connectivity in the rodent. Brain Connect. 2013;3(1):31–40. doi:10.1089/brain.2012.0115

46. Thompson GJ, Pan WJ, Keilholz SD. Different dynamic resting state fMRI patterns are linked to different frequencies of neural activity. J Neurophysiol. Jul 2015;114(1):114–24. doi:10.1152/jn.00235.2015

47. Abbas A, Belloy M, Kashyap A, et al. Quasi-periodic patterns contribute to functional connectivity in the brain. Neuroimage. May 1 2019;191:193–204. doi:10.1016/j.neuroimage.2019.01.076

48. Ribatti D, Nico B, Crivellato E. The role of pericytes in angiogenesis. Int J Dev Biol. 2011;55(3):261–8. doi:10.1387/ijdb.103167dr

49. Salhia B, Angelov L, Roncari L, Wu X, Shannon P, Guha A. Expression of vascular endothelial growth factor by reactive astrocytes and associated neoangiogenesis. Brain Res. Nov 10 2000;883(1):87–97. doi:10.1016/s0006-8993(00)02825-0

50. Bonkowski D, Katyshev V, Balabanov RD, Borisov A, Dore-Duffy P. The CNS microvascular pericyte: pericyte-astrocyte crosstalk in the regulation of tissue survival. Fluids Barriers CNS. Jan 18 2011;8(1):8. doi:10.1186/2045-8118-8-8

51. Bogaert E, Van Damme P, Poesen K, et al. VEGF protects motor neurons against excitotoxicity by upregulation of GluR2. Neurobiol Aging. Dec 2010;31(12):2185–91. doi:10.1016/j.neurobiolaging.2008.12.007

52. Ellison SM, Trabalza A, Tisato V, et al. Dose-dependent neuroprotection of VEGF(1)(6)(5) in Huntington’s disease striatum. Mol Ther. Oct 2013;21(10):1862–75. doi:10.1038/mt.2013.132

53. Storkebaum E, Carmeliet P. VEGF: a critical player in neurodegeneration. J Clin Invest. Jan 2004;113(1):14–8. doi:10.1172/JCI20682

54. Liu H, Zhang C, Xu J, et al. Huntingtin silencing delays onset and slows progression of Huntington’s disease: a biomarker study. Brain. Nov 29 2021;144(10):3101–3113. doi:10.1093/brain/awab190

55. Whitesell JD, Liska A, Coletta L, et al. Regional, Layer, and Cell-Type-Specific Connectivity of the Mouse Default Mode Network. Neuron. Feb 3 2021;109(3):545–559 e8. doi:10.1016/j.neuron.2020.11.011

56. Donzis EJ, Estrada-Sanchez AM, Indersmitten T, et al. Cortical Network Dynamics Is Altered in Mouse Models of Huntington’s Disease. Cereb Cortex. Apr 14 2020;30(4):2372–2388. doi:10.1093/cercor/bhz245

57. Indersmitten T, Tran CH, Cepeda C, Levine MS. Altered excitatory and inhibitory inputs to striatal medium-sized spiny neurons and cortical pyramidal neurons in the Q175 mouse model of Huntington’s disease. J Neurophysiol. Apr 1 2015;113(7):2953–66. doi:10.1152/jn.01056.2014

58. Wang Y, Sepers MD, Xiao D, Raymond LA, Murphy TH. Water-Reaching Platform for Longitudinal Assessment of Cortical Activity and Fine Motor Coordination Defects in a Huntington Disease Mouse Model. eNeuro. Jan 2023;10(1)doi:10.1523/ENEURO.0452-22.2022

59. Margulies DS, Ghosh SS, Goulas A, et al. Situating the default-mode network along a principal gradient of macroscale cortical organization. Proc Natl Acad Sci U S A. Nov 1 2016;113(44):12574–12579. doi:10.1073/pnas.1608282113

60. Vazquez-Rodriguez B, Suarez LE, Markello RD, et al. Gradients of structure-function tethering across neocortex. Proc Natl Acad Sci U S A. Oct 15 2019;116(42):21219–21227. doi:10.1073/pnas.1903403116

61. Huntenburg JM, Yeow LY, Mandino F, Grandjean J. Gradients of functional connectivity in the mouse cortex reflect neocortical evolution. Neuroimage. Jan 15 2021;225:117528. doi:10.1016/j.neuroimage.2020.117528

62. Shafiei G, Baillet S, Misic B. Human electromagnetic and haemodynamic networks systematically converge in unimodal cortex and diverge in transmodal cortex. PLoS Biol. Aug 2022;20(8):e3001735. doi:10.1371/journal.pbio.3001735

63. Fulcher BD, Murray JD, Zerbi V, Wang XJ. Multimodal gradients across mouse cortex. Proc Natl Acad Sci U S A. Mar 5 2019;116(10):4689–4695. doi:10.1073/pnas.1814144116

64. Chuang KH, Li Z, Huang HH, Khorasani Gerdekoohi S, Athwal D. Hemodynamic transient and functional connectivity follow structural connectivity and cell type over the brain hierarchy. Proc Natl Acad Sci U S A. Jan 31 2023;120(5):e2202435120. doi:10.1073/pnas.2202435120

65. Zarkali A, McColgan P, Leyland LA, Lees AJ, Rees G, Weil RS. Organisational and neuromodulatory underpinnings of structural-functional connectivity decoupling in patients with Parkinson’s disease. Commun Biol. Jan 19 2021;4(1):86. doi:10.1038/s42003-020-01622-9

66. Glasser MF, Van Essen DC. Mapping human cortical areas in vivo based on myelin content as revealed by T1- and T2-weighted MRI. J Neurosci. Aug 10 2011;31(32):11597–616. doi:10.1523/JNEUROSCI.2180-11.2011

67. Kubera KM, Schmitgen MM, Hirjak D, Wolf RC, Orth M. Cortical neurodevelopment in pre-manifest Huntington’s disease. Neuroimage Clin. 2019;23:101913. doi:10.1016/j.nicl.2019.101913

68. Barnat M, Capizzi M, Aparicio E, et al. Huntington’s disease alters human neurodevelopment. Science. Aug 14 2020;369(6505):787–793. doi:10.1126/science.aax3338

69. van der Plas E, Schultz JL, Nopoulos PC. The Neurodevelopmental Hypothesis of Huntington’s Disease. J Huntingtons Dis. 2020;9(3):217–229. doi:10.3233/JHD-200394

70. van der Plas E, Langbehn DR, Conrad AL, et al. Abnormal brain development in child and adolescent carriers of mutant huntingtin. Neurology. Sep 3 2019;93(10):e1021–e1030. doi:10.1212/WNL.0000000000008066

71. Mehler MF, Gokhan S. Mechanisms underlying neural cell death in neurodegenerative diseases: alterations of a developmentally-mediated cellular rheostat. Trends Neurosci. Dec 2000;23(12):599–605. doi:10.1016/s0166-2236(00)01705-7

72. Cepeda C, Oikonomou KD, Cummings D, et al. Developmental origins of cortical hyperexcitability in Huntington’s disease: Review and new observations. J Neurosci Res. Dec 2019;97(12):1624–1635. doi:10.1002/jnr.24503

73. Braz BY, Wennagel D, Ratie L, et al. Treating early postnatal circuit defect delays Huntington’s disease onset and pathology in mice. Science. Sep 23 2022;377(6613):eabq5011. doi:10.1126/science.abq5011

74. Zhang C, Wu Q, Liu H, et al. Abnormal Brain Development in Huntington’ Disease Is Recapitulated in the zQ175 Knock-In Mouse Model. Cereb Cortex Commun. 2020;1(1):tgaa044. doi:10.1093/texcom/tgaa044

75. Tereshchenko AV, Schultz JL, Bruss JE, Magnotta VA, Epping EA, Nopoulos PC. Abnormal development of cerebellar-striatal circuitry in Huntington disease. Neurology. May 5 2020;94(18):e1908–e1915. doi:10.1212/WNL.0000000000009364

76. Khakh BS, Beaumont V, Cachope R, Munoz-Sanjuan I, Goldman SA, Grantyn R. Unravelling and Exploiting Astrocyte Dysfunction in Huntington’s Disease. Trends Neurosci. Jul 2017;40(7):422–437. doi:10.1016/j.tins.2017.05.002

77. Khakh BS, Goldman SA. Astrocytic contributions to Huntington’s disease pathophysiology. Ann N Y Acad Sci. Mar 2 2023;doi:10.1111/nyas.14977

78. Jansen AH, van Hal M, Op den Kelder IC, et al. Frequency of nuclear mutant huntingtin inclusion formation in neurons and glia is cell-type-specific. Glia. Jan 2017;65(1):50–61. doi:10.1002/glia.23050

79. Tong X, Ao Y, Faas GC, et al. Astrocyte Kir4.1 ion channel deficits contribute to neuronal dysfunction in Huntington’s disease model mice. Nat Neurosci. May 2014;17(5):694–703. doi:10.1038/nn.3691

80. Diaz-Castro B, Gangwani MR, Yu X, Coppola G, Khakh BS. Astrocyte molecular signatures in Huntington’s disease. Sci Transl Med. Oct 16 2019;11(514)doi:10.1126/scitranslmed.aaw8546

81. Hall CN, Reynell C, Gesslein B, et al. Capillary pericytes regulate cerebral blood flow in health and disease. Nature. Apr 3 2014;508(7494):55–60. doi:10.1038/nature13165

82. Brown LS, Foster CG, Courtney JM, King NE, Howells DW, Sutherland BA. Pericytes and Neurovascular Function in the Healthy and Diseased Brain. Front Cell Neurosci. 2019;13:282. doi:10.3389/fncel.2019.00282

83. Hsiao HY, Chen YC, Huang CH, et al. Aberrant astrocytes impair vascular reactivity in Huntington disease. Ann Neurol. Aug 2015;78(2):178–92. doi:10.1002/ana.24428

84. Drouin-Ouellet J, Sawiak SJ, Cisbani G, et al. Cerebrovascular and blood-brain barrier impairments in Huntington’s disease: Potential implications for its pathophysiology. Ann Neurol. Aug 2015;78(2):160–77. doi:10.1002/ana.24406

85. Lange C, Storkebaum E, de Almodovar CR, Dewerchin M, Carmeliet P. Vascular endothelial growth factor: a neurovascular target in neurological diseases. Nat Rev Neurol. Aug 2016;12(8):439–54. doi:10.1038/nrneurol.2016.88

86. Ruiz de Almodovar C, Lambrechts D, Mazzone M, Carmeliet P. Role and therapeutic potential of VEGF in the nervous system. Physiol Rev. Apr 2009;89(2):607–48. doi:10.1152/physrev.00031.2008

87. Cepeda C, Levine MS. Synaptic Dysfunction in Huntington’s Disease: Lessons from Genetic Animal Models. Neuroscientist. Feb 2022;28(1):20–40. doi:10.1177/1073858420972662

88. Foster NN, Barry J, Korobkova L, et al. The mouse cortico-basal ganglia-thalamic network. Nature. Oct 2021;598(7879):188–194. doi:10.1038/s41586-021-03993-3

89. Hintiryan H, Foster NN, Bowman I, et al. The mouse cortico-striatal projectome. Nat Neurosci. Aug 2016;19(8):1100–14. doi:10.1038/nn.4332

90. Smith EJ, Sathasivam K, Landles C, et al. Early detection of exon 1 huntingtin aggregation in zQ175 brains by molecular and histological approaches. Brain Commun. 2023;5(1):fcad010. doi:10.1093/braincomms/fcad010

