## Supplementary information file for "Evolution of aberrant brain-wide spatiotemporal dynamics of resting-state networks in a Huntington’s disease mouse model"

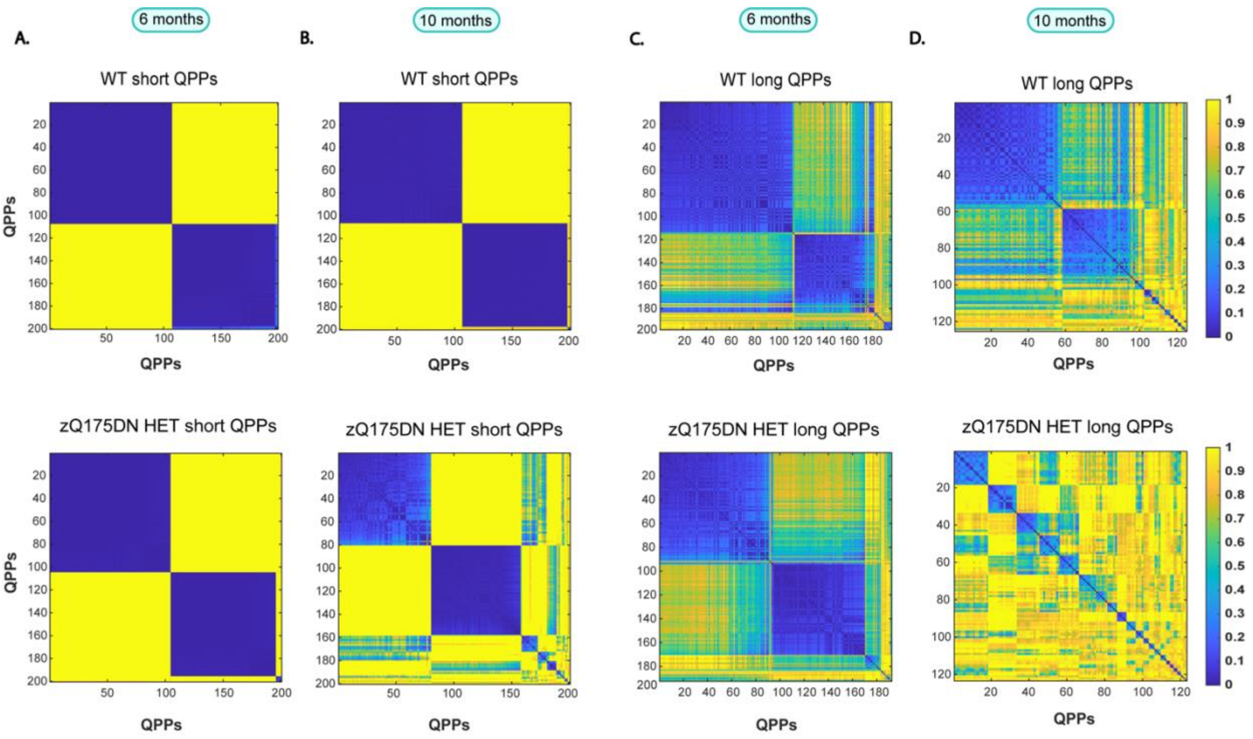


**<u>Figure S1.</u> Spatiotemporal hierarchical clustering of short (3s) and long (10s) QPPs in WT and zQ175DN HET mice at 6 and 10 months.** Matrices representing (dis)similarity of QPPs obtained in WT (top) and zQ175DN HET (bottom) for short (A, B) and long (C, D) QPPs; blue – yellow color bar represents the closest to most distant QPPs based on their spatiotemporal properties

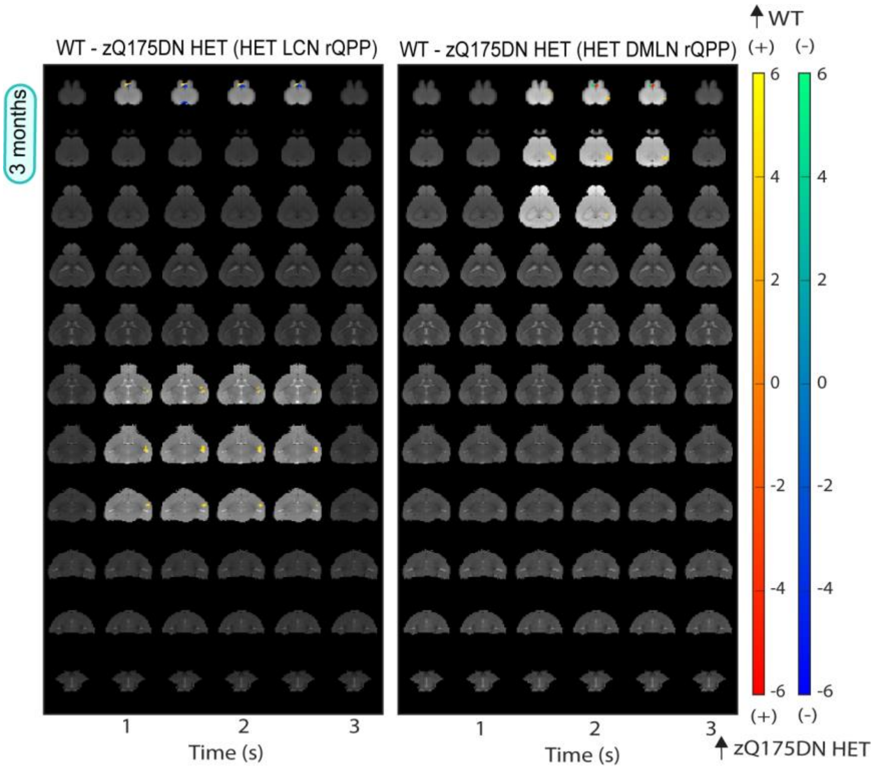


**<u>Figure S2.</u> Spatial activation differences in HET rLCN and rDMLN QPP at 3 months of age.** Two-sample T-test maps (two-tailed, FDR, $p < 0.05$) of between-group differences where the red-yellow colour bar displays differences in positively activated whereas the blue-green colour bar represents activation differences in negatively activated voxels.

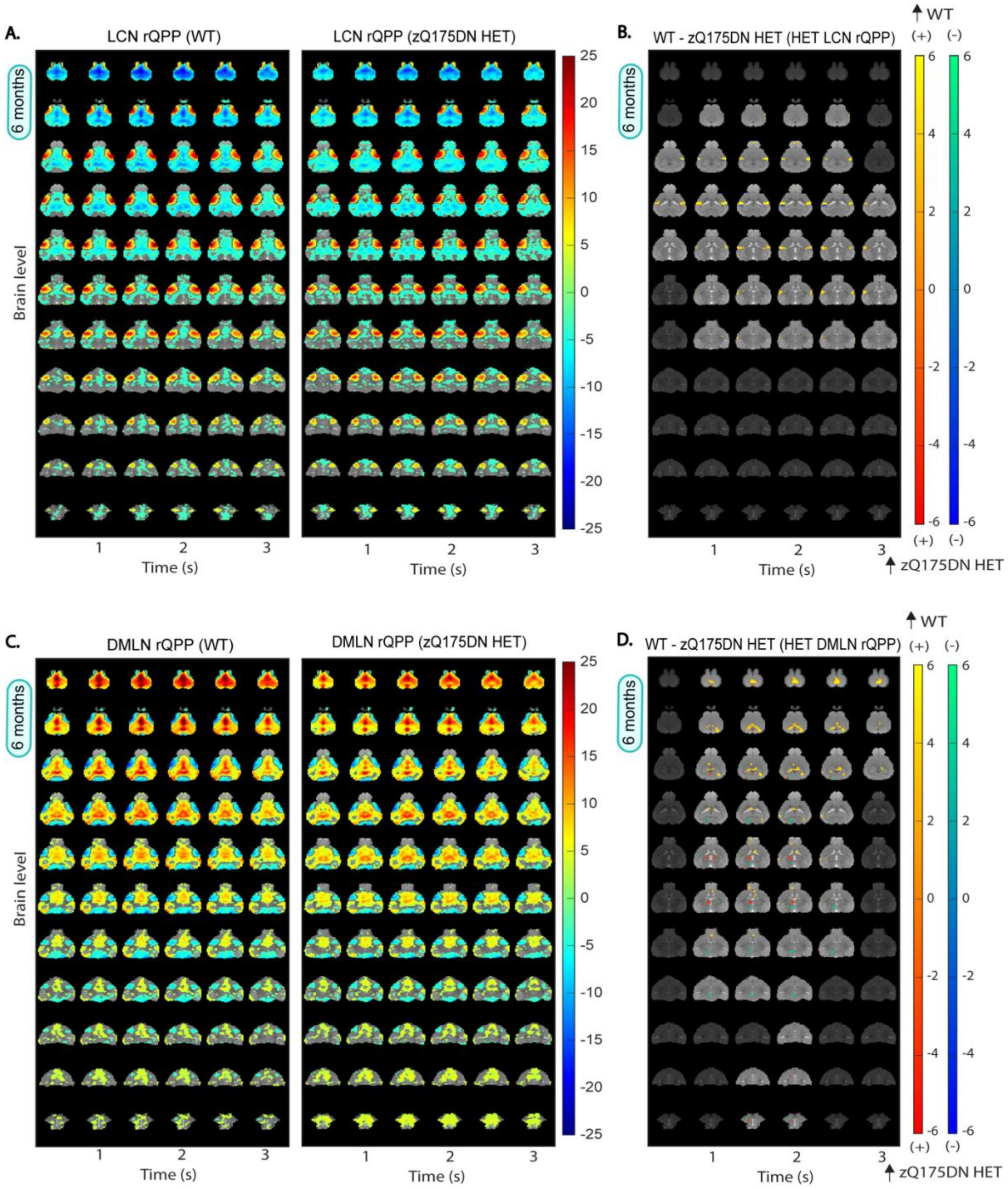


<u>Figure S3.</u> **Group representative DMLN and LCN QPPs in WT and zQ175DN HET and spatial activation differences in zQ175DN rQPPs at 6 months of age.** (A, C) One-sample T-maps of rLCN and rDMLN QPPs in both groups where significantly activated (red-yellow) and deactivated (green-blue) voxels as compared to the mean BOLD signal are shown (two-tailed one-sample T-test, FDR, $p < 0.05$) (B, D) Two-sample T-test maps (two-tailed, FDR, $p < 0.05$) of between-group differences shown for the zQ175DN HET rLCN and rDMLN QPP; Red-yellow colour bar displays differences in positively activated whereas the blue-green colour bar represents activation differences in negatively activated voxels.

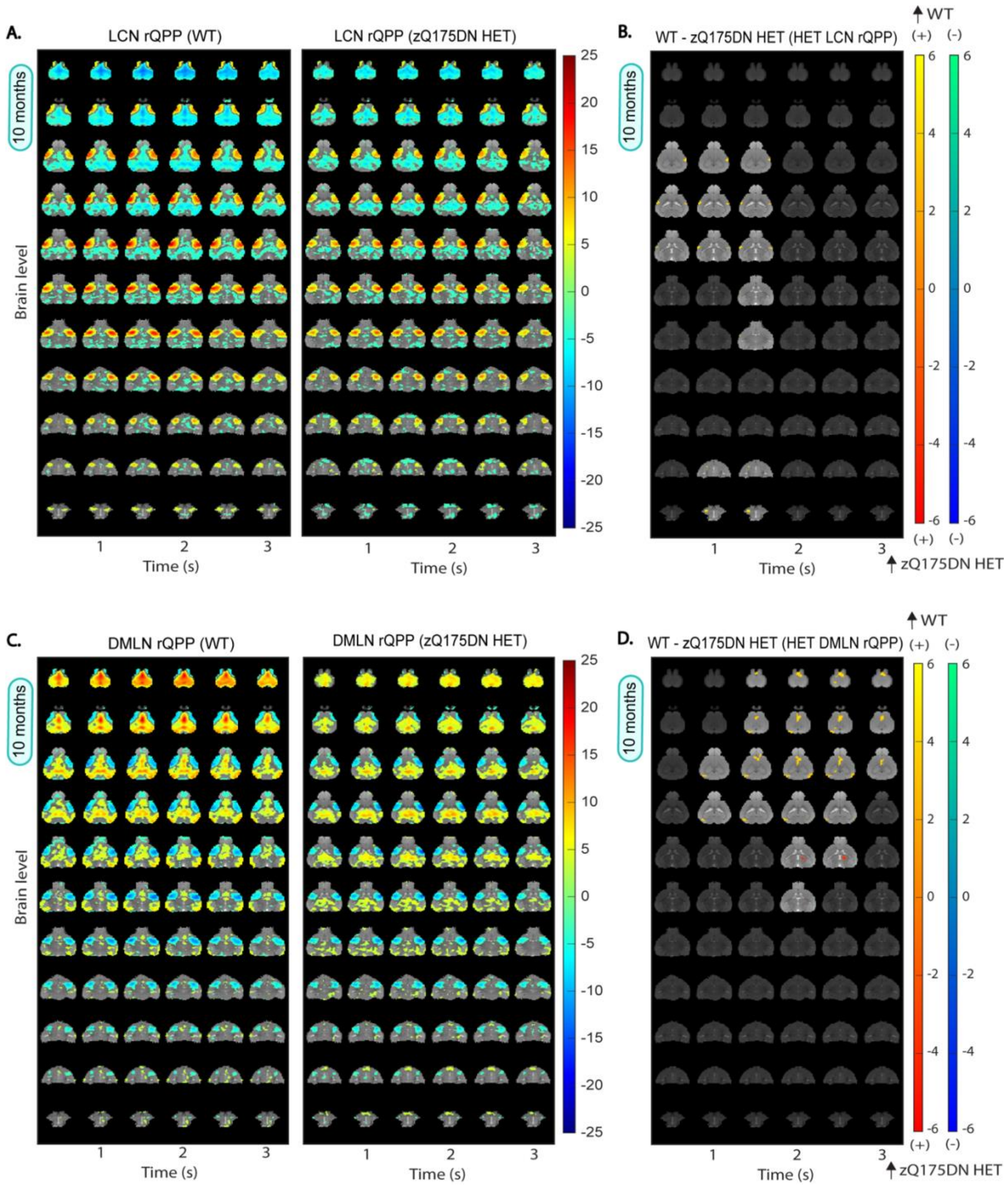


**Figure S4.** **Group representative DMLN and LCN QPPs in WT and zQ175DN HET and spatial activation differences in zQ175DN rQPPs at 10 months of age.** (A, C) One-sample T-maps of rLCN and rDMLN QPPs in both groups where significantly activated (red-yellow) and deactivated (green-blue) voxels as compared to the mean BOLD signal are shown (two-tailed one-sample T-test, FDR, $p < 0.05$) (B, D) Two-sample T-test maps (two-tailed, FDR, $p < 0.05$) of between-group differences shown for the zQ175DN HET rLCN and rDMLN QPP; Red-yellow colour bar displays differences in positively activated whereas the blue-green colour bar represents activation differences in negatively activated voxels.

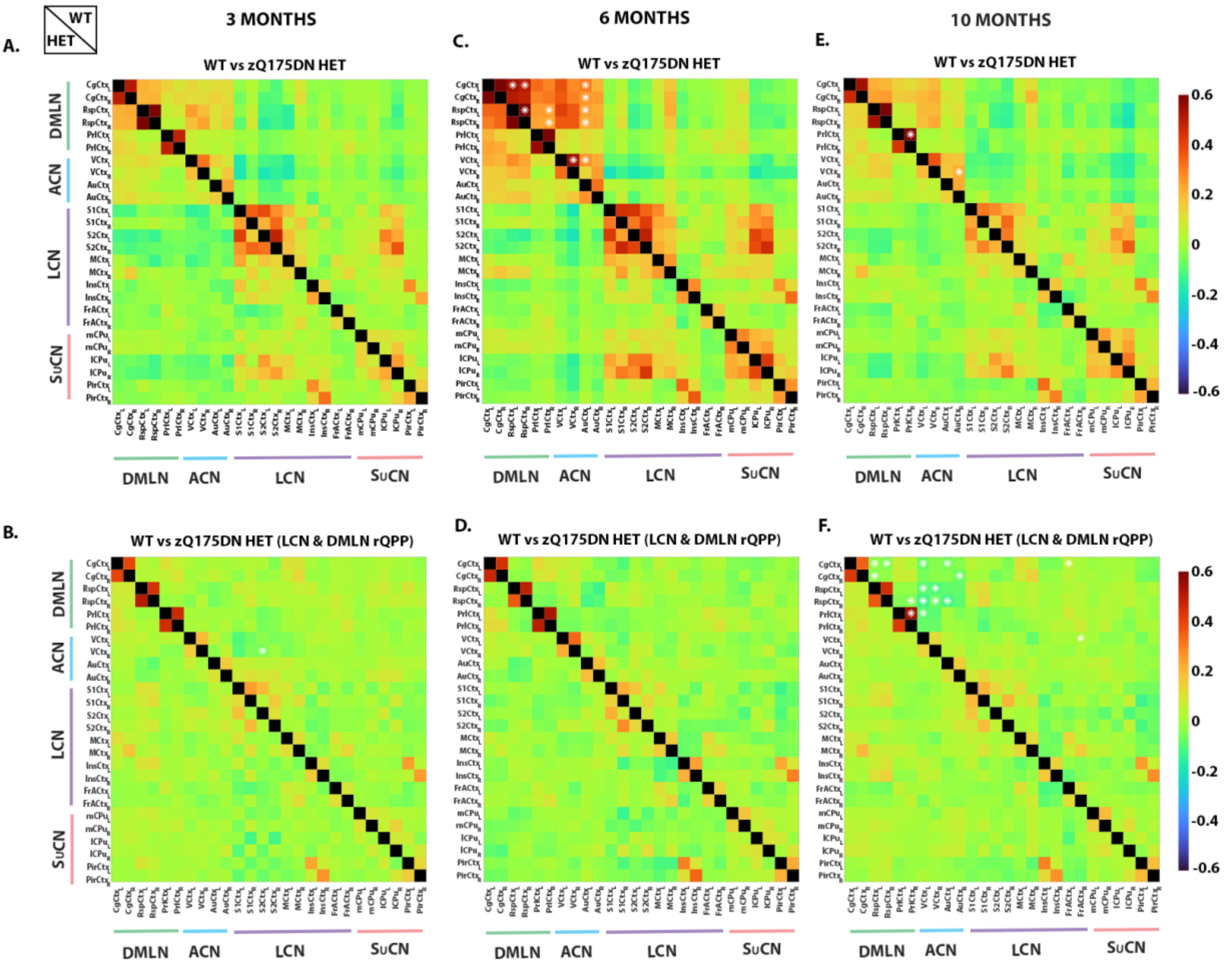


**<u>Figure S5.</u> Regression of LCN and DMLN rQPPs and their contribution to FC genotypic differences across different ages.** Mean z-transformed correlation of FC in WT (top) and zQ175DN HET (bottom) matrices of each group (A, C, E) and FC after regression of LCN and DMLN rQPPs in each group (B, D, F) at 3, 6 and 10 months of age; red/orange colours represent positively correlated connectivity, green color indicates very low to no connectivity between regions and dark/light blue colours represent negatively correlated connectivity; White asterisks (top) indicate significant group differences in FC based on a two-sample T-test ($p < 0.05$, FDR) only performed on connections that demonstrated a significant FC in at least one group based on a one-sample T-test.

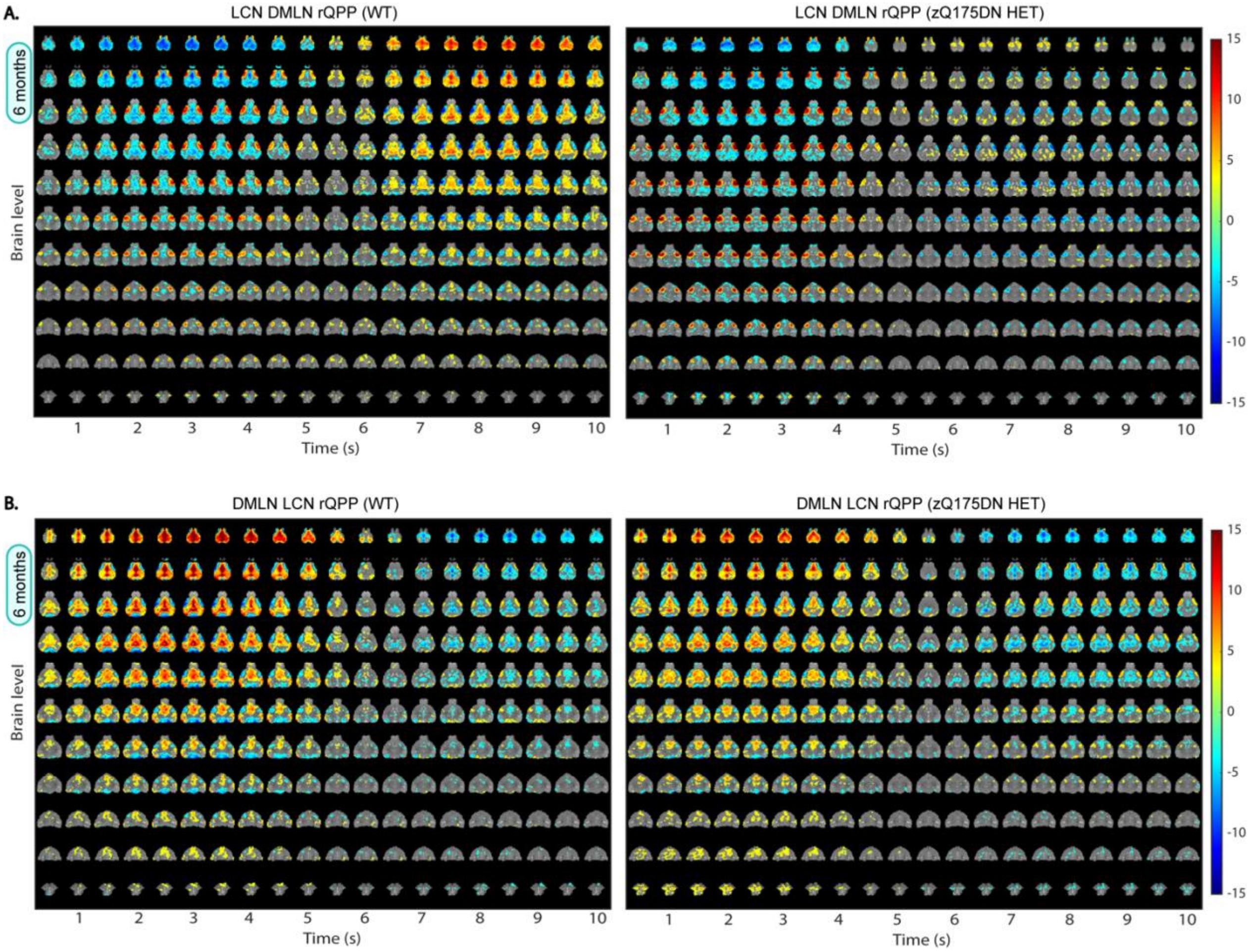


**<u>Figure S6.</u> Group rLCN DMLN and rDMLN LCN QPPs in WT and zQ175DN HET at 6 months of age.** One-sample T-maps of (A) rLCN DMLN QPP and (B) rDMLN LCN QPP in both groups where significantly activated (red-yellow) and deactivated (green-blue) voxels as compared to the mean BOLD signal are shown (two-tailed one-sample T-test, FDR, $p < 0.05$).

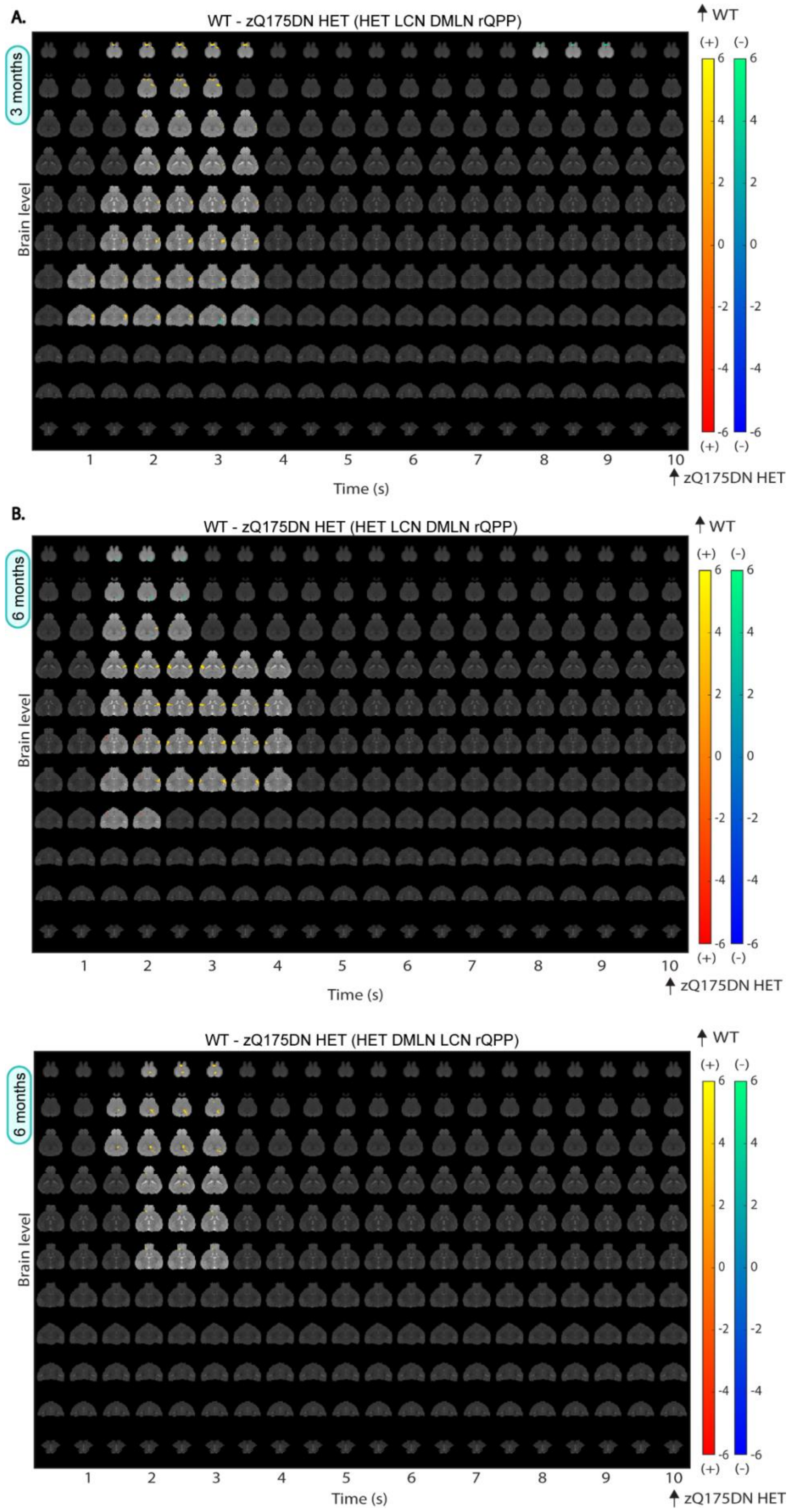
A.
WT - zQ175DN HET (HET LCN DMLN rQPP)
3 months
Brain level
Time (s)
WT
zQ175DN HET
(+)
(-)
B.
WT - zQ175DN HET (HET LCN DMLN rQPP)
6 months
Brain level
Time (s)
WT
zQ175DN HET
WT - zQ175DN HET (HET DMLN LCN rQPP)
6 months
Brain level
Time (s)
WT
zQ175DN HET

**<u>Figure S7.</u> Spatial activation differences in HET rLCN DMLN and rDMLN LCN QPP at 3 and 6 months of age.** Two-sample T-test maps (two-tailed, FDR, $p < 0.05$) of between-group differences where the red-yellow colour bar displays differences in positively activated whereas the blue-green colour bar represents activation differences in negatively activated voxels.

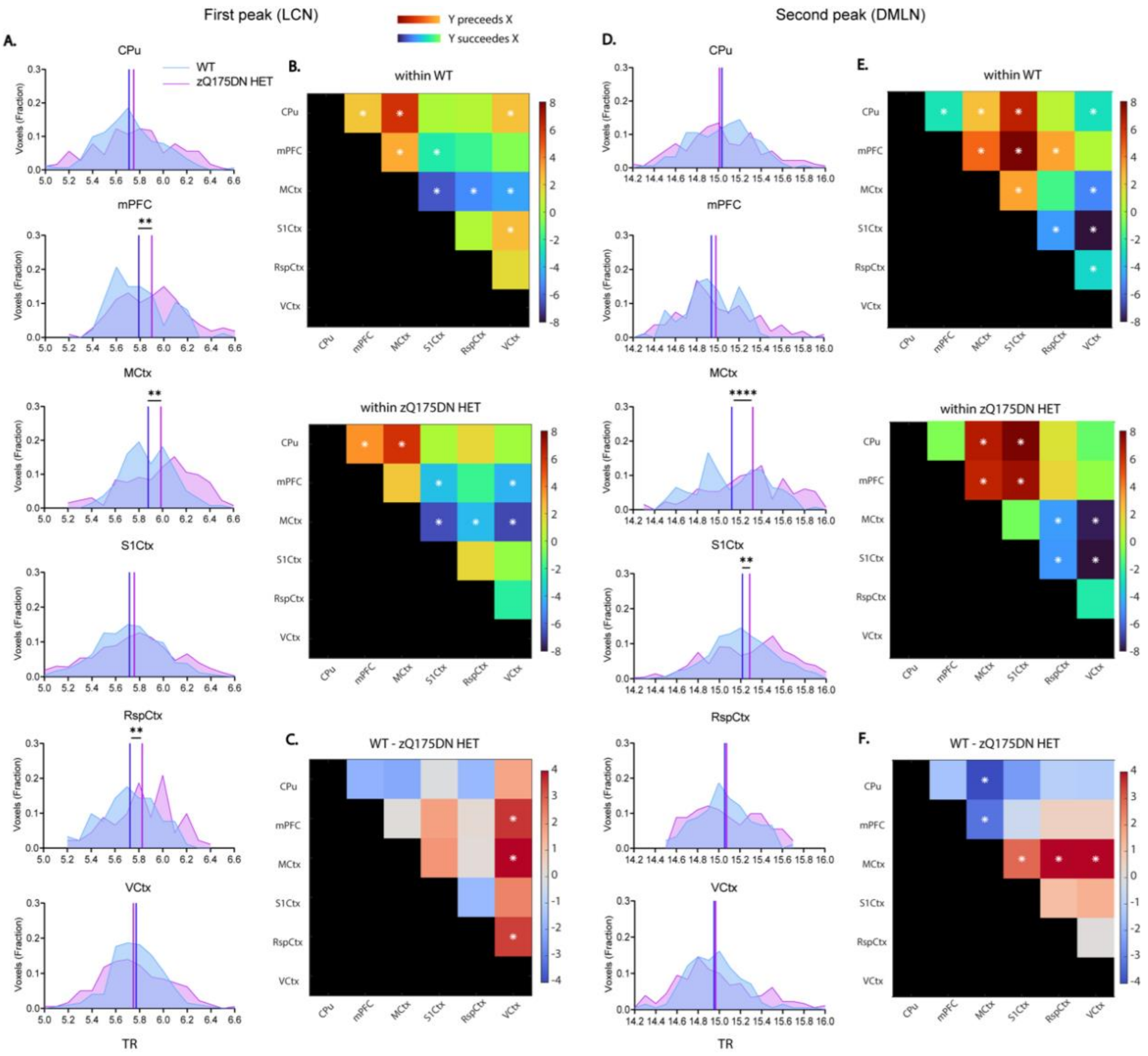


**Figure S8.** **Regional mean timing and inter-regional timing differences in WT LCN DMLN rQPP at 3 months of age.** (A, D) histogram of the distribution of significant voxels with the between-group differences in mean peak timings for each region during the first (LCN) and second (DMLN) peak of WT LCN DMLN rQPP (B, E) between region timing relationship within each group; (C, F) between-group differences of inter-regional timing intervals; All comparisons were corrected for both metrics and all regions (two-sample t-test, FDR, $p < 0.05$); CPu caudate putamen, mPFC medial prefrontal cortex, MCtx motor cortex, SS1Ctx Somatosensory Cortex 1, RspCtx retrosplenial cortex, VCtx visual cortex;

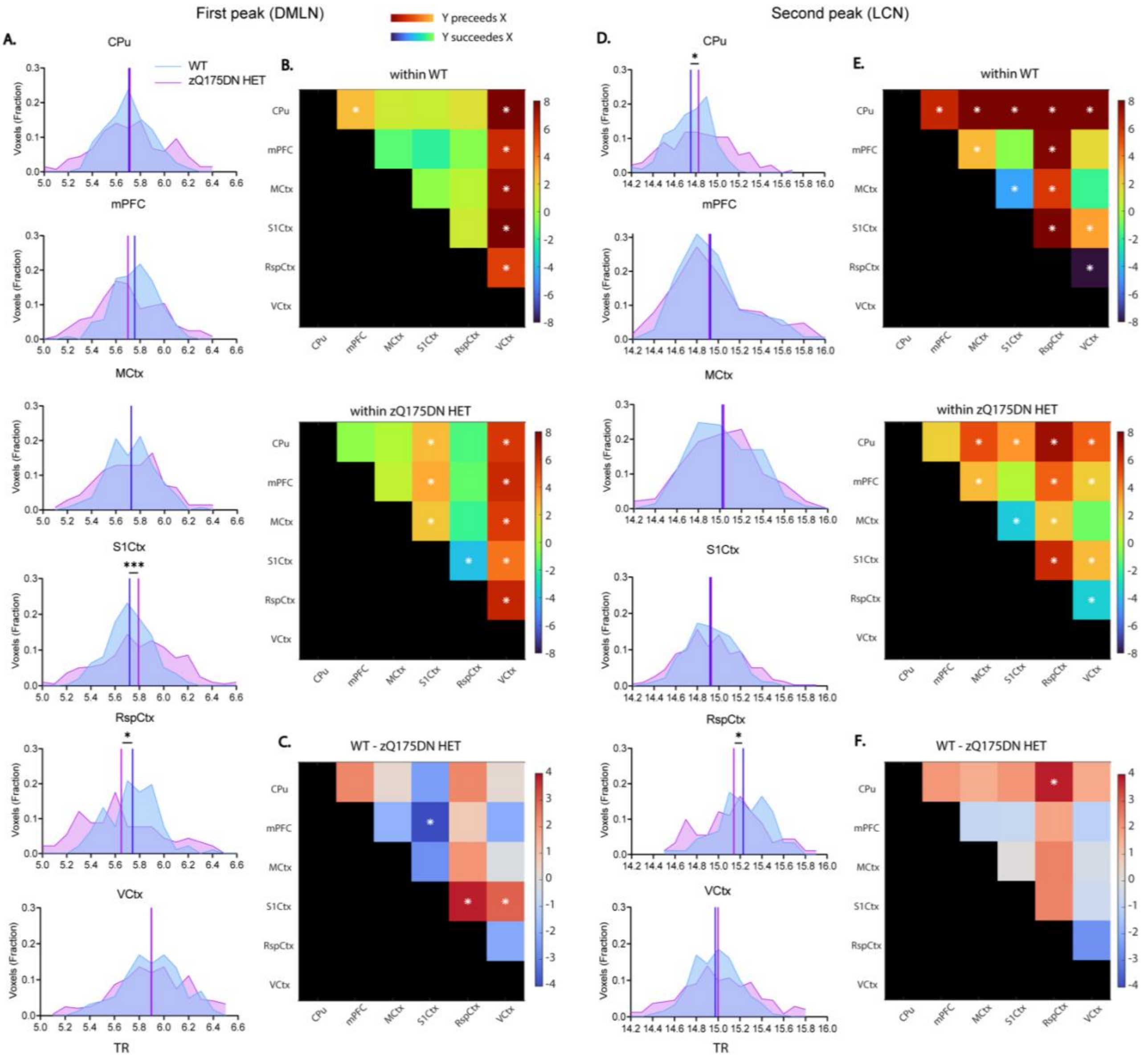


**Figure S9. Regional mean timing and inter-regional timing differences in WT DMLN LCN rQPP at 6 months of age.** (A, D) histogram of the distribution of significant voxels with the between-group differences in mean peak timings for each region during the first (LCN) and second (DMLN) peak of WT LCN DMLN rQPP (B, E) between region timing relationship within each group; (C, F) between-group differences of inter-regional timing intervals; All comparisons were corrected for both metrics and all regions (two-sample t-test, FDR, $p < 0.05$); CPu caudate putamen, mPFC medial prefrontal cortex, MCtx motor cortex, SS1Ctx Somatosensory Cortex 1, RspCtx retrosplenial cortex, VCtx visual cortex;

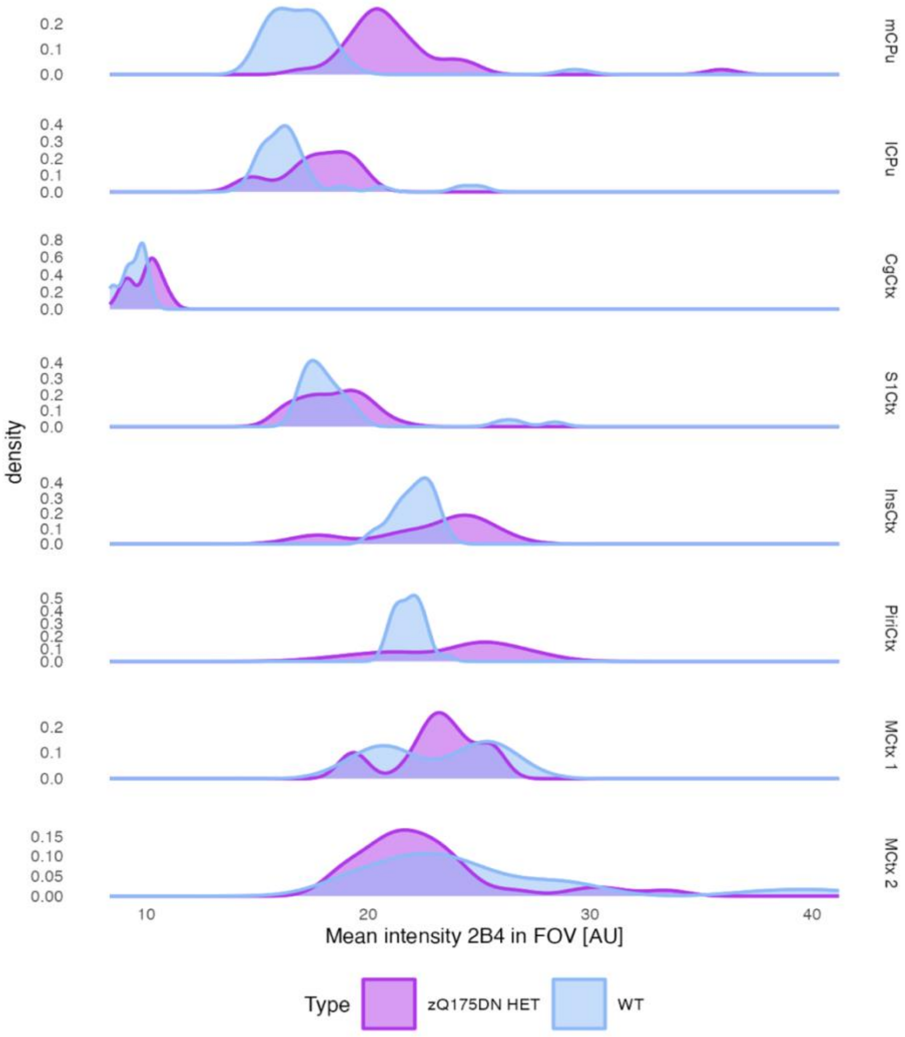


**<u>Figure S10.</u> Distribution of the mean 2B4 intensities per FOV across different regions in WT and zQ175DN HET at 8 months of age.** mCPu medial caudate putamen, lCPu lateral caudate putamen, CgCtx cingulate cortex, S1Ctx somatosensory cortex 1, InsCtx insular cortex, PiriCtx piriform cortex, MCtx 1 motor cortex 1, MCtx 2 motor cortex 2.

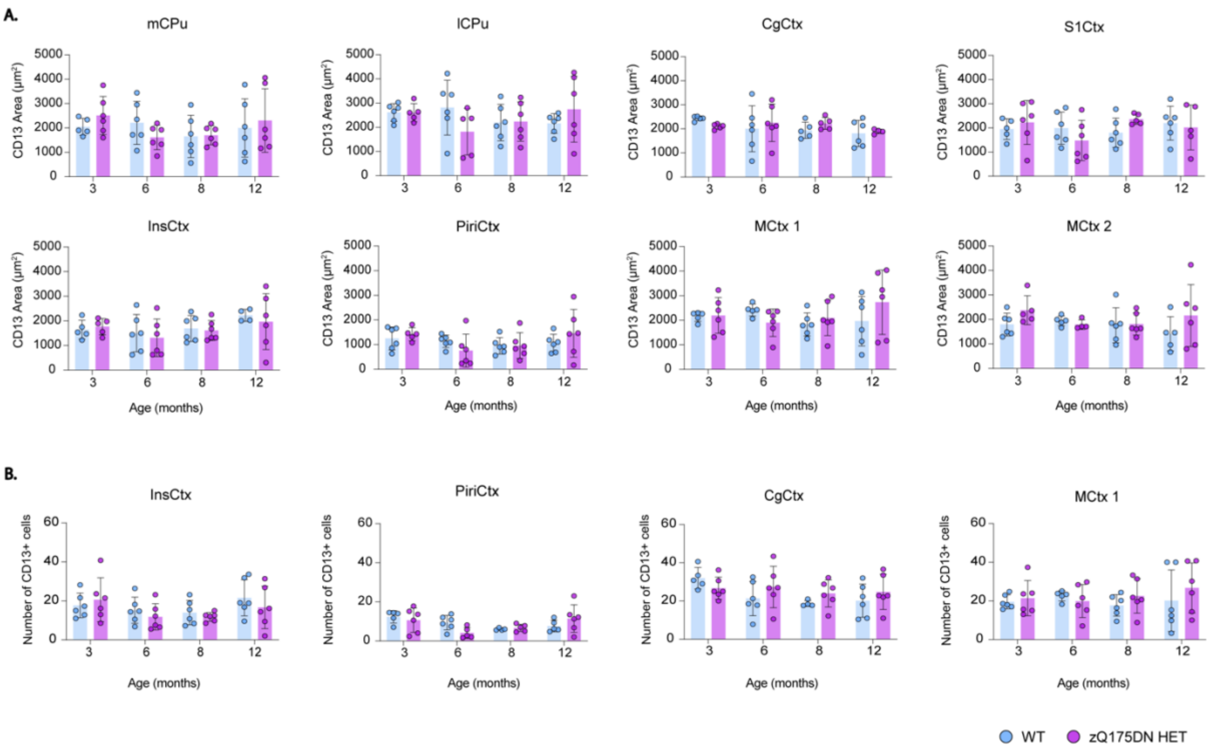


**<u>Figure S11.</u> Area, and count of pericytes in the zQ175DN and WT mice across ages.** between-group comparison of (A) Total CD13+ area relative to FOV across ages (B) number of CD13+ cells across ages; (mean ± SD, two-way ANOVA, $p < 0.05$); mCPu medial caudate putamen, lCPu lateral caudate putamen, PiriCtx piriform cortex, InsCtx insular cortex, MCtx 1 motor cortex 1, MCtx 2 motor cortex 2, S1Ctx somatosensory cortex 1, CgCtx cingulate cortex.

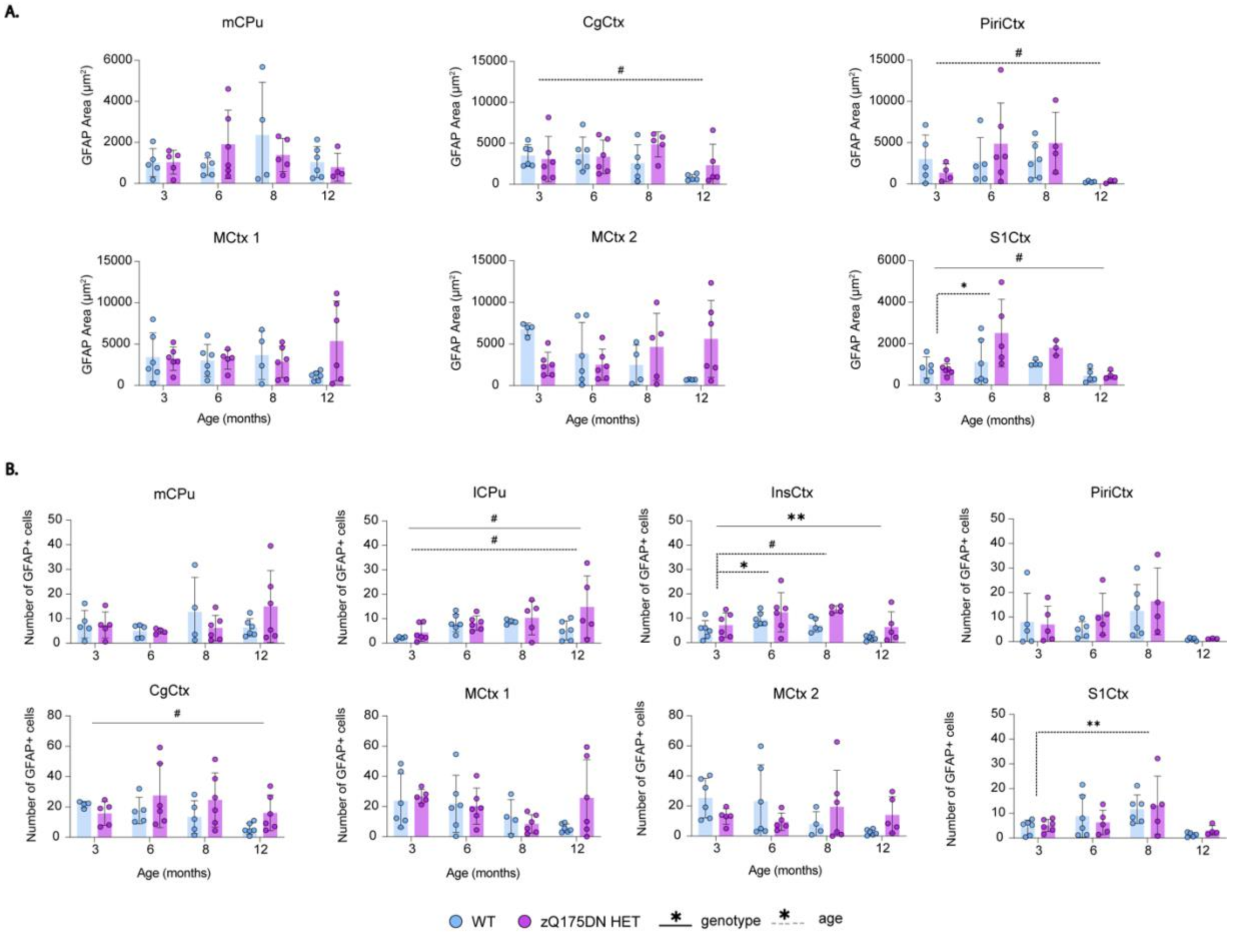


**Figure S12.** **Area and count of astrocytes in the zQ175DN HET and WT mice across ages.** (A) thresholded GFAP+ area relative to FOV and (B) number of GFAP+ cells (mean ± SD, two-way ANOVA, $p < 0.05$); * $p \leq 0.05$, ** $p \leq 0.01$, *** $p \leq 0.001$, **** $p \leq 0.0001$, # $p \leq 0.1$; mCPu medial caudate putamen, lCPu lateral caudate putamen, PiriCtx piriform cortex, InsCtx insular cortex, MCtx 1 motor cortex 1, MCtx 2 motor cortex 2, S1Ctx somatosensory cortex 1, CgCtx cingulate cortex;
